# Life history traits and dispersal shape neutral genetic diversity in metapopulations

**DOI:** 10.1101/2021.07.13.452195

**Authors:** Jimmy Garnier, Pierre Lafontaine

## Abstract

Genetic diversity at population scale, depends on species life-history traits, population dynamics and local and global environmental factors. We first investigate the effect of life-history traits on the neutral genetic diversity of a single population using a deterministic mathematical model. When the population is stable, we show that semelparous species with precocious maturation and iteroparous species with delayed maturation exhibit higher diversity because their life history traits tend to balance the lifetimes of non reproductive individuals (juveniles) and adults which reproduce. Then, we extend our model to a metapopulation to investigate the additional effect of dispersal on diversity. We show that dispersal may truly modify the local effect of life history on diversity. As a result, the diversity at the global scale of the metapopulation differ from the local diversity which is only described through local life history traits of the populations. In particular, dispersal usually promotes diversity at the global metapopulation scale.

## 1 Introduction

Environmental changes induced by climate changes or human activities, disrupt population and metapopulation dynamics, resulting in species extinctions, profound changes in ecosystem dynamics or loss of genetic variation (Ceballos and Ehrlich, 2002; Haddad et al., 2015). Through its relation with demographic processes (Mittell et al., 2015; Vilas et al., 2015), neutral genetic diversity is often used to inform about the evolutionary and demographic history of populations (Paz-Vinas et al.,2018). At population scale, the genetic diversity depends on species life-history traits, population dynamics and local and global environmental factors (Eckert et al., 2008; Attard et al., 2015). Even though many theoretical and empirical studies have shown the importance of local population dynamics in shaping species’ range or communities (e.g.Husband and Barrett, 1996; Freckleton and Watkinson, 1990; Levin et al., 2003), few is known about the relative importance of local and range wide processes in driving genetic diversity. Moreover, understanding the relationship between neutral genetic diversity, species life history traits and environmental factors might be a key to establish general conservation guidelines valid across taxa and space (Willoughby et al., 2015; Blanchet et al., 2017).

Plants and animals life histories exhibit a huge diversity. Individuals may live for hours or for centuries; they may reproduce once in their lifetime (semelparity) or several time once they become reproductive adult (iteroparity). Individuals may experiment several totally different niches during their lifetime. Specialized stages may exist for dispersal or dormancy. Moreover, the life history traits of individuals, such as survival probabilities, development, fecundity or dispersal, almost always depend on their age, their body size or their developmental stage. The resulting variety of life history strategies has profound consequences on genetic diversity (Nelson et al., 2005; Bonnefon et al., 2013). However, the studies only focus on the effect of one particular life history trait or consider them independently whereas genetic diversity patterns more likely result from the interaction between life-history traits and potentially environmental factors. For instance, empirical studies have found that populations of small animals with high fecundity and short longevity, large geographic ranges and long-distance dispersal harbour relatively high genetic diversity (Eo et al.,2011; Romiguier et al., 2014; Doyle et al., 2015; Dalongeville et al., 2016), whereas other studies could not validate this life-history related genetic diversity pattern (Mitton and Lewis, 1989; Vachon et al., 2018).

Despite the lack of studies investigating the combined effect of life-history traits and environmental factors on genetic diversity at the local scale of the population, various quantitative reviews focused on both life-history traits and spatial factors to understand genetic diversity across populations that is at the global scale of species or metapopulation (Schoville et al., 2012; Romiguier et al.,2014; Miraldo et al., 2016; Manel et al., 2020). In particular, these studies demonstrated that the genetic diversity is lower for long-lived or low-fecundity species than for short-lived or high-fecundity species. However, these studies focusing on genetic diversity at species scale do not capture the local-scale processes ruled for instance by environmental constraints or anthropogenic factors. Conversely to genetic diversity at species or metapopulation scale, the genetic diversity at population scale is driven by the population dynamics reflecting the local ability of the population to cope with environmental conditions. In addition, although species genetic diversity provides crucial insights into species’ demographic trajectories, it is poorly informative in terms of contemporary population dynamics or on the risk of genetic erosion of populations. Thus, it is important to provide a framework to access the impact of life-history traits and environmental conditions on the genetic diversity at the population and metapopulation scale.

Here, we provide a mathematical framework that incorporates the two genetic diversity scales and makes the link between population dynamics and neutral genetic diversity. First, we use the stage-classified demography framework to incorporate the diversity of life histories into population models. And we combine this approach with classical metapopulation models to describe the variety of environmental conditions or life-history strategies among species range. Our resulting matrix model allows us to describe the stage-structure in each local population (Leslie, 1945; Lefkovitch,1965) as well as the dispersal between those populations. On top of that, we use the mathematical tool introduced by Garnier et al. (2012) and Roques et al. (2012) to study the spatio– temporal dynamics of the neutral genetic diversity in a range–expanding population. Their framework, inspired from a simulation study of Hallatschek and Nelson (2008), had already been applied to a wide class of reaction–dispersion model (Bonnefon et al., 2014). More recently, they extend their work to metapopulation models described through a system of ordinary differential equations in a fully heterogeneous environment (Garnier and Lafontaine, 2021). In the present paper, we extend the inside dynamics approach which links the demography dynamics of the metapopulation and the neutral genetic diversity, to our matrix projection model.

More precisely, we consider a metapopulation of genes or haploid individuals composed of local panmictic populations structured by stage and living in different habitat patches linked by dispersal. Our stage–structured metapopulation model describes the population density **N**(*t*) = (**N**_1_(*t*)),…, **N**_*ω_h_*_(*t*) including the vector of population densities **N**_*k*_(*t*) in each habitat *k* at time *t*, over *ω_h_* habitats. Each vector of population densities **N**_*k*_(*t*) = (*N*_1,*k*_(*t*),…, *N_ω_c_,k_*(*t*) includes the population densities *N_i,k_*(*t*) of individuals of stage *i* living in habitat *k*. We assume that the number of stages *ω_c_* is the same for any habitat. Our structured metapopulation model takes the following form, for each habitat *k*

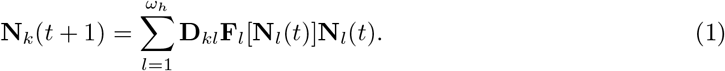

It indicates that the projection interval is divided into two main phases of possible different duration: reproduction phase (**F**) followed by a dispersal phase (**D**). The reproduction matrices **F**_*k*_ describe the life cycle dynamics of the local stage–structured population living in the habitat *k* (Neubert and Caswell, 2000). They depend on the life-history traits of the population and the environmental conditions of the habitat. The dispersal matrices 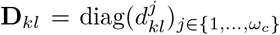 describe the proportion of individuals of stage *j* that moves from habitat *l* to habitat *k* during one time step. They depend on both the arrival habitat *k* and the departure habitat *l* and they may also depend on the stage j of the individuals which are moving.

In this paper, we first describe the dynamics of neutral alleles when the gene dynamics follows the general stage–structured metapopulation (1). Then, we focus on a single population structured by two stages: reproducing adults and non-reproducing juveniles. This simple situation permits to explore the impact of four life history traits (survival of juveniles and adults, development rate between juvenile and adult, and fecundity) on the genetic diversity in a population at steady state (equilibrium or periodic steady state). Finally, we explore the intertwined effect of these life-history traits, the dispersal and environmental heterogeneity on genetic diversity of a two-stage structured metapopulation located over two distinct habitats. In particular, we will be able to understand the effect of the juvenile stage on the neutral genetic diversity of a metapopulation. As already observed in literature, the presence of juvenile stage helps to promote genetic diversity in range expanding population (Bonnefon et al., 2013; Marculis et al., 2019) as well as predator-prey systems (Nelson et al., 2005). Moreover, we aim to assess the impact of life history of species through reproduction strategy (semelparous or iteroparous strategies) and development strategy (delayed or precocious development) on the neutral genetic diversity.

## 2 Materials and methods

### 2.1 Allele dynamics and diversity indices

Our structured metapopulation model assumes discrete and overlapping generations and we neglect mutation and random genetic drift.

We consider a single locus with *I* neutral alleles. Thus the genes or haploid individuals in the metapopulation are characterized by their alleles. For each allele *i* ∈ {1,…, *I*}, its density is represented by the matrix 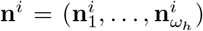, where the vectors 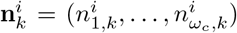 represent the allele densities in habitat *k*, and 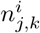 represent the allele densities in the habitat *k* among genes carried by individuals of stage *j*. Thus, in each habitat *k*, the population density **N**_*k*_ = (*N*_1_,_*k*_,…, *N*_*ω_c_*,*k*_) in the metapopulation **N** satisfying (1), is equal to the sum of the allele densities in habitat *k* (the allele density might be zero if this allele is absent of the habitat). More precisely, for each stage *j* in habitat *k* we have:

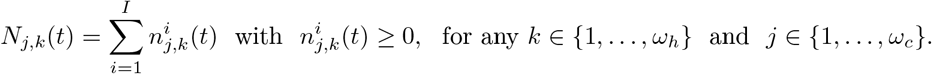

In addition, we assume that the alleles are neutral so the genes (or the haploid individuals) only differ by their initial location, their initial stage and their alleles. In particular, they share the same dispersal ability. Moreover, in each habitat the genes (or the haploid individuals) follow the same reproduction characteristics as the other genes with the same stage. More precisely, in each habitat *k* the allele density 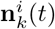 evolves according to the reproduction matrix **F**_*k*_[**N**_*k*_(*t*)] in this habitat. In addition, the migration ability of each allele **n**^*i*^ is described by the dispersal matrix **D** = (**D**_*kl*_)_*k,l*∈{1,…,*ω_h_*_} of the metapopulation. Each allele density **n**^*i*^(*t*) therefore satisfies the following linear dynamical system:

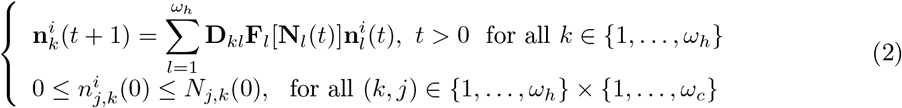

where 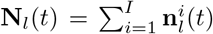 for any habitat *l* ∈ {1,…,*ω_h_*}. Throughout the remaining sections, we use the superscript *i* to denote the neutral alleles and the subscript *j* for the stage and the subscript *k* for the habitat. Note that the number of neutral alleles *I* does not need to be equal to *ω_h_* × *ω_c_* the product of the number of stages *ω_c_* and the number of habitats *ω_h_*.

This decomposition method provides a mathematical framework to describe and analyze the neutral genetic diversity dynamics of our stage–structured metapopulation. More precisely, for each allele *i* ∈ {1,…,*I*}, we can define its frequency at different scale: *p^i^* at metapopulation scale, 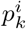 in each habitat *k* and 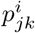 in each habitat *k* and each stage *j*. Thus, we can define the neutral genetic diversity at different scales: the metapopulation scale (*γ*–diversity), the habitat scale or the stage scale (*α*–diversity). The neutral *γ*–diversity, corresponding to the total neutral genetic diversity in the metapopulation is quantified through the following index *γ-Div*(*t*):

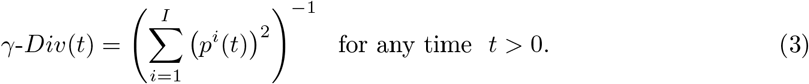

This diversity index corresponds to the inverse of the Simpson index (Simpson, 1949) which describes the probability that two individuals sampled randomly in habitat *k* among stages at time *t* belong to the same allele *i*. It is also the inverse of the total homozygosity in the metapopulation. A high index of diversity indicates high diversity or a true evenness in the population: *γ-Div* is maximal when all the alleles frequencies are equal, i.e., when *p*^1^ = … = *p*^I^ = 1/*I*.

At the scale of habitat, we describe the neutral *α_h_*–diversity corresponding to the harmonic mean neutral genetic diversity within habitats by the harmonic mean of local diversity indices *γ-Div_k_*(*t*) in habitat *k* weighted by the proportion *P_k_* of the metapopulation living in habitat *k*:

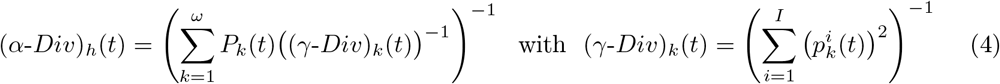

This index is also the inverse of the mean homozygosity across habitats.

Similarly, at the stage scale, we can also define the neutral *α_c_*–diversity corresponding to the harmonic mean neutral genetic diversity among habitat within a stage by the harmonic mean of local stage diversity indices *γ-Div_jk_*(*t*) in habitat *k* and stage *j* weighted by the proportion *P_jk_* of the metapopulation of stage *j* living in habitat *k*:

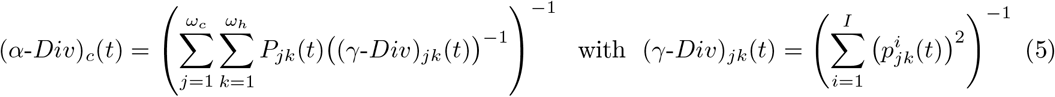

This index is also the inverse of the mean homozygosity across stages.

### 2.2 Dispersal and demographic model

#### Dispersal matrix D

The dispersal matrix **D** is a block matrix where each square matrix **D**_*kl*_ is diagonal 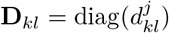 and the dispersal rate 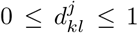 may depend on the stage *j* ∈ {1,…, *ω_c_*} of the individuals moving from habitat *k* to habitat *l*. Moreover, we have the following relationship between the dispersal matrices 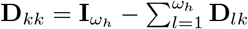 to ensure that the matrix **D** is a dispersal matrix.

For instance, if we consider a metapopulation over two habitats and structured with two stages (juveniles and adults), we write:

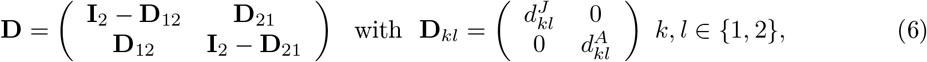

where 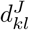 and 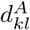 are the dispersal rate from habitat *l* to habitat *k* of respectively the juveniles and the adults.

#### Reproduction matrix F

The reproduction matrix or projection matrix **F** = diag(**F**_*k*_)_*k*∈{1…*ω_h_*}_ describes in each habitat *k*, the reproduction, survival and the interactions between stages in this habitat. So for each habitat *k*, the reproduction matrix **F**_*k*_ only depends on the population **N**_*k*_ in this habitat *k* and its entries should be non negative:

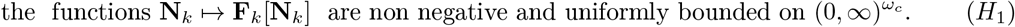

Moreover, we assume that density-dependence occurs during life-cycle. Thus, the reproduction matrix **F**_*k*_ in each habitat *k* depends on the population density in this habitat **N**_*k*_. Moreover, the spectral radius of those matrices *ρ*(**F**_*k*_[**N**_*k*_]) satisfies

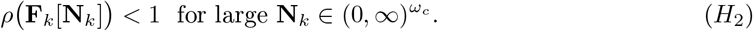

For instance, if we assume that our metapopulation is structured with only two stages (*ω_c_* = 2), the juveniles *J* and the adults *A* (**N**_*k*_ = (*J_k_*, *A_k_*) for any *k* ∈ {1,…, *ω_h_*}), then the following projection matrix for each habitat *k* satisfies the hypotheses above:

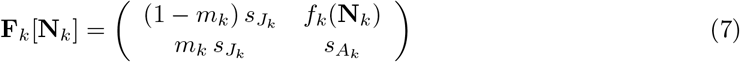

where *s _J_k__* and are the survival probability of respectively the juveniles and the adults, *m_k_* is the maturation probability and *f_k_* is the fecundity which depends on the population **N**_*k*_ = (*J_k_*, *A_k_*) in habitat *k, f_k_*(**N**_*k*_) = f_0*k*_ exp (–(*J_k_* + *A_k_*)/*β_k_*) where *f*_0*k*_ is the intrinsic fecundity and *β_k_* quantifies the density dependence of the fecundity with respect to the population density.

#### Survival and steady states

First, let us remark that equation (1) can take the following form, where the metapopulation density matrix **N** is written as a vector **N** = ((*N*_1,1_…, *N_ω_c__*, 1),…, (*N*_1,*ω_h_*_,…, *N_ω_c_,ω_h__*)):

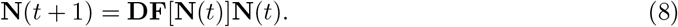

Moreover, we assume that the metapopulation never goes extinct over the different habitats even if some habitats are not favourable. More precisely, we assume that the spectral radius of the matrix **DF**[**0**] satisfies

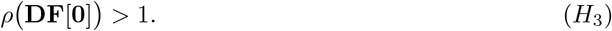

This hypothesis (*H*_3_) ensures that the steady state **0** is unstable. Moreover, we know from fixed point theorem that a non-negative steady state **N**\* exists for our model and it satisfies:

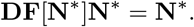

In order to ensure the existence of positive steady state, we assume the following hypothesis,

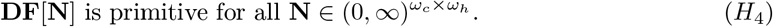

This hypothesis implies in particular that the dispersal matrices **D_*kl*_** cannot be identically equal to **0** for all *l* ∈ {1,…, *ω_h_*}, otherwise, the matrix **DF**[**N**] is reducible. Under the hypothesis (*H*_4_), the steady states **N**\* of (1) are positive thanks to the Perron-Frobenius theorem. In particular, for the example above with only 1 habitat, we can compute the unique positive steady state **N**\* = (*J*\*, *A*\*) as follows:

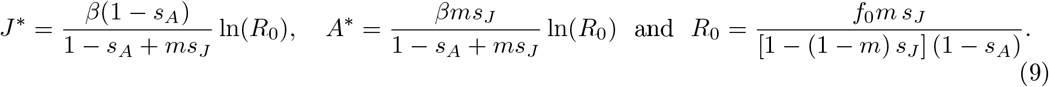

This equilibrium is stable until some parameters, such that fecundity *f*_0_ or dispersal 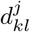, reach critical values where stability is lost. In this case, bifurcations occur which may generate different dynamics: cycles, quasi-cycles or even chaos (Neubert and Caswell, 2000). The cycles are characterized by positive periodic steady states **N**^*^(*t*) (see bifurcation diagram Fig. 1(a) and the Appendix A for more details). This phenomenon may also occur when the metapopulation is composed of several habitats (see bifurcation diagram Fig. 1(b) for the case of 2 habitats).

**Figure 1:**
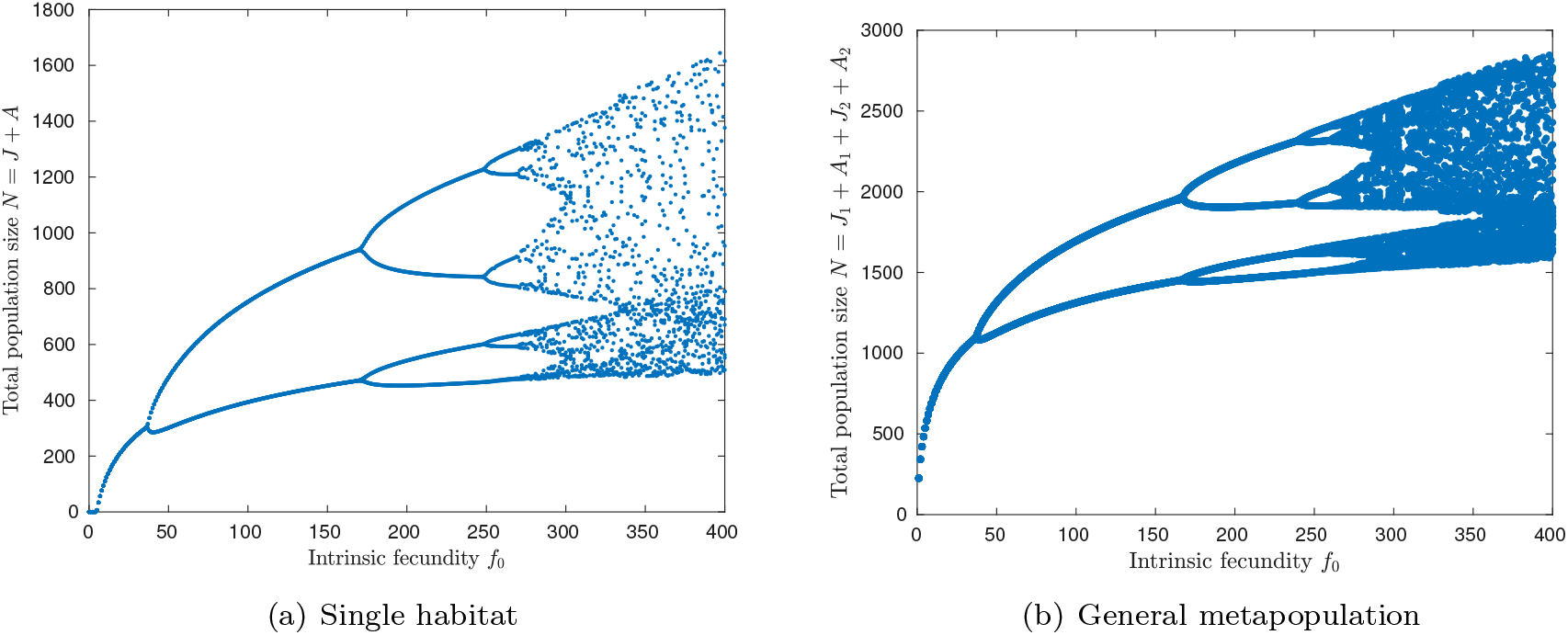
Bifurcation diagram of model (1) with respect to the intrinsic fecundity *f*_0_ for a single population (a) and a metapopulation composed of two habitats (b). The fecundity ranges in (0, 400). The characteristics of habitats are *β*_1_ = *β*_2_ = 150, *s*_*J*_1__ = *s*_*j*_2__ = 0.5, *s*_*A*_1__ = 0.2, *m*_1_ = 0.2 and *s*_*A*_2__ = 0.9, *m*_2_ = 0.9. And the dispersal rate is 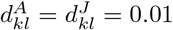.

## 3 Allele dynamics in a stage–structured metapopulation

First, we aim to describe the dynamics of the neutral alleles in the stage–structured metapopulation. Then, we investigate the effect of the life-history traits (adult survival, maturation rate, dispersal) on the neutral genetic diversity. In the following section we always assume that hypotheses (*H*_1_)-(*H*_4_) are satisfied to ensure the existence of positive steady states **N**\*.

### 3.1 Neutral allele dynamics and allele frequencies in a metapopulation at steady state

Henceforth, we assume that the metapopulation is at steady state, that is **N**(*t*) = **N**^*^(*t*) where **N**\* is either a stationary state of (1) or a *T*–periodic steady state of (1). We investigate the dynamics of a particular neutral allele **n** in this metapopulation which is described by the following equation:

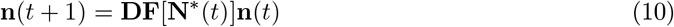

starting from **n**(0) = (*n*_1,1_(0)…, *n_ω_c_,1_(0)),…, (*n*_1,*ω*_h__*(0),…, *n*_ω_c_,*ω_h_*_(0))) such that 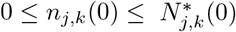 for all (*j, k*) ∈ {1,…, *ω_c_*} × {1,…,*ω_h_*}. We have the following result.

#### Theorem 1.

*Let* **n**(*t*) *be the solution of*(10) *starting from* **n**(0) *such that* 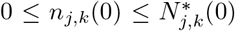 *for all* (*j*, *k*) ∈ {1,…, *ω**c***} × {1,…, *ω_h_*}, *then*:

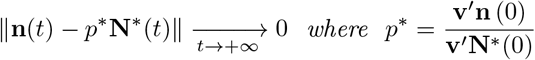

*and* **v**′ *is the transposed eigenvector of either the transpose of the matrix* **DF**[**N**\*] *associated to the main eigenvalue* 1 *if* **N**\* *is a stationary state or the transpose of the matrix* 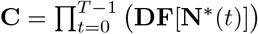 *if* **N**\* *is a T–periodic steady state*.

Our result, proved in section 7.1, describes the dynamics of the alleles in the metapopulation at any stage level (see Fig. 2(a)) and for various steady states of the metapopulation (see Fig. 3). It also provides analytical expression of the asymptotic frequency *p*^*^ of any neutral allele in the metapopulation. As expected from classical population genetics model this asymptotic frequency depends neither on the location nor on the stage of the individuals. However, the asymptotic allele frequency differs from the initial allele frequency (see Fig. 2(a)). Indeed in a single habitat, we can start with 3 alleles with the same frequency in the population (see Fig. 2(b)) but different frequency among each stage (see Fig. 2(b)) then the allele frequencies will eventually converge toward different values in the population (see Fig. 2(a)). Although the population is at demographic equilibrium, the allele frequencies evolve through time and their evolution depends on the spatial structure of the metapopulation through the dispersal as well as the stage structure through the reproduction (see paragraph above for more mathematical insights).

**Figure 2:**
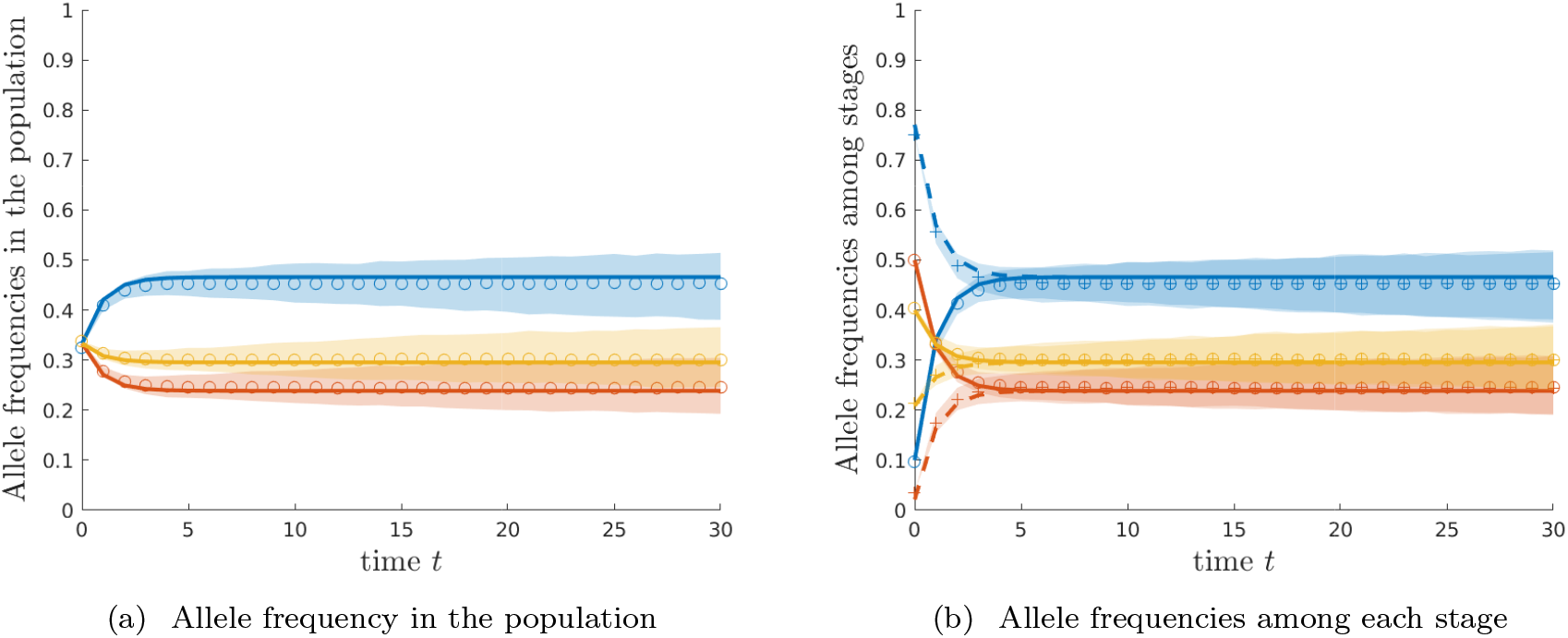
Temporal dynamics of 3 alleles in a single habitat for the deterministic model (plain and dashed lines) and the IBM model for 5.10^3^ individuals (circle and cross marked curves correspond tothe medians and the shaded regions correspond to the 99% confidence intervals over 10^3^ replicates).Each colour corresponds to one allele: in panel (a) the allele frequency in the population (juvenilesand adults) and in panel (b), the allele frequency among each stage (plain curve corresponds tojuveniles and dashed curves to adults). Habitat characteristics: *f*_01_ = 1.5, *s_A_1__* = 0.7, *s*_*J*_1__ = 0.8, *m*_1_ = 0.2, *β*_1_ = 100 and *N_s_* = 50.

**Figure 3:**
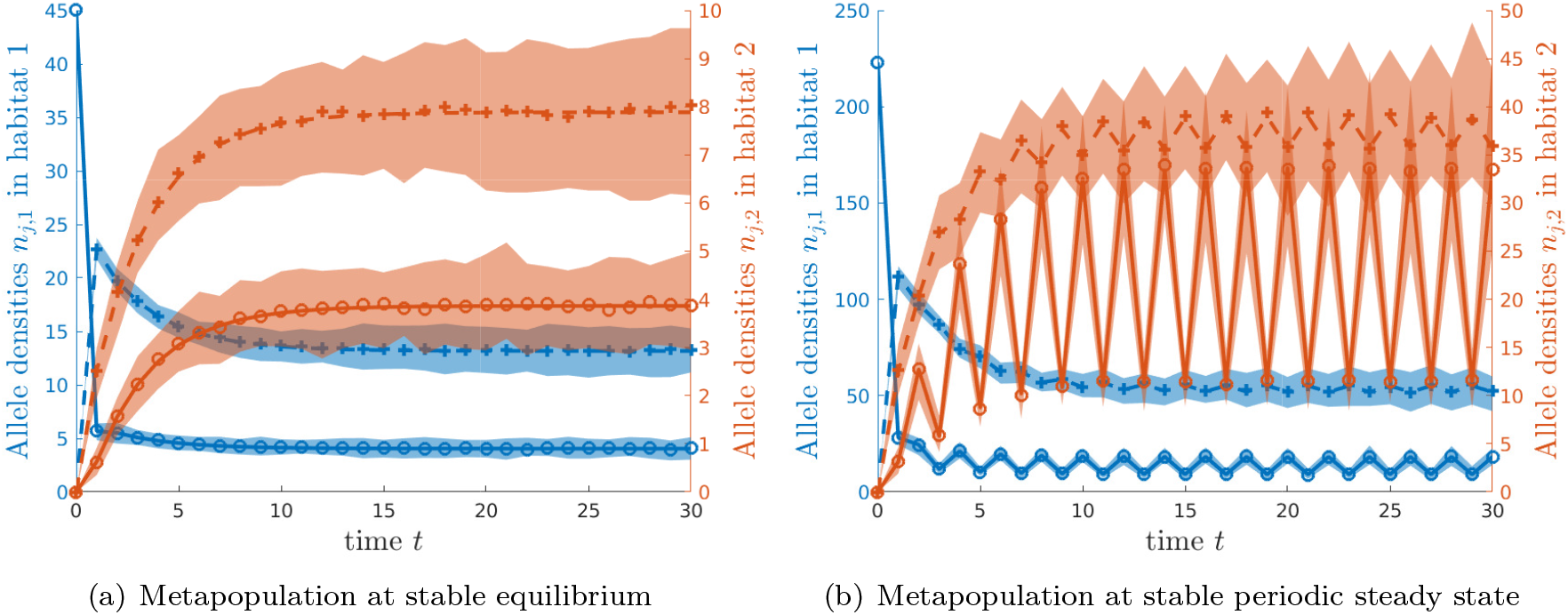
Temporal dynamics of one neutral allele in a stage-structure metapopulation composed of 2 connected habitats and 2 stages (juvenile and adult): (a) at equilibrium (small fecundity rate *f*_01_ = *f*_02_ = 1.5) and (b) at periodic steady state (large fecundity rate *f*_01_ = *f*_02_ = 450) for the deterministic model (plain and dashed lines) and the IBM model for 5.10^3^ individuals (circle and star marked curves correspond to the medians and the shaded regions correspond to the 99% confidence intervals over 10^3^ replicates). Blue curves correspond to the densities in habitat 1 and red curves in habitat 2 (plain curve corresponds to juveniles and dashed curves to adults). In habitat 1: *s*_*A*_1__ = 0.8, *s*_J_1__ = 0.7, *m*_1_ = 0.8, *β*_1_ = 150, *ε*_12_ = 0.1 and in habitat 2: *s*_*A*2_ = 0.9, *s*_*J*_2__ = 0.6, *m*_2_ = 0.5, *β*_2_ = 150 and *ε*_21_ = 0.2

Moreover, thanks to the primitivity property of the projection matrix **DF**[**N**\*], we know that **v** is also positive and thus *p*\* is positive. So in this situation, any allele initially present in the metapopulation will persist where it is initially present. Moreover, it will eventually spread over the entire habitats and it will generate individuals of any stages. This result shows that the richness of genetic alleles is preserved globally (any allele initially present persists for ever in the metapopulation). Moreover, the local richness either in each habitat or in each stage among the metapopulation may be enhanced thanks to the demography dynamics. This result stresses that allele frequency are not always fixed in the metapopulation over time. If an allele is absent from a habitat, then initially its frequency is 0 while it eventually converges toward *p*\* > 0.

However, if the projection matrix **DF**[**N**\*] is reducible (no more primitive nor irreducible), some alleles may go extinct which reduces the global genetic richness of the metapopulation. Indeed, let us consider a metapopulation composed of two habitats in which individuals can only move from habitat 2 to habitat 1, that is **D**_12_ = **0** in the definition of **D** stated in (6). Then, any alleles that are initially only in habitat 1 will go extinct, that is *p*^*^ = 0 from theorem 1 (see Appendix B.1 for mathematical details). Thus dispersal should have important impact on neutral genetic diversity.

#### Analytical insights for the two-stage isolated populations at stationary state

In the general situation, it is not easy to express the eigenvector **v** with respect to the equilibrium **N**^*^. So for further insights, let us look at a simple case where all the populations are isolated and they are not connected with each other, that is **D** is the identity matrix. In addition, we assume that the populations are structured with two stages (juveniles and adults) and the reproduction matrix **F**_*k*_ in each habitat *k* is given by equation (7). Since all the populations are isolated we can just look at one population. In this population, we can compute explicitly the asymptotic proportion *p*^*^ of any neutral allele **n** initially located in this population with density **n**(0):

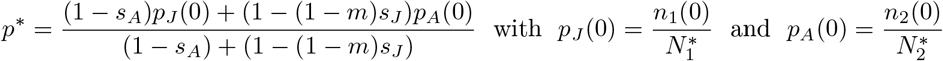

where *p_J_*(0) and *p_A_*(0) represent respectively the initial allele frequency among juveniles, respectively adults, in the population **n** (see equation (15) for details). First, we notice that the asymptotic frequency *p*\* of both stages truly differ from their initial proportions *p_J_*(0) and *p_A_*(0). So, the demographic dynamics truly shapes the neutral allele frequencies in the population. For instance, if initially the proportions in each stage of the allele are identical, that is *p_J_*(0) = *p_A_*(0), then the asymptotic allele frequency remains the initial allele frequency. However, if initially the allele frequencies in each stage are different, then they will evolve in time towards the asymptotic allele frequency *p*\* (see Fig. 2), which is also different than the initial allele frequency among the population defined by

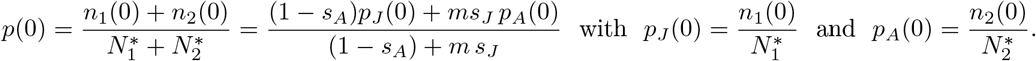

The stage–structure of the population has a profound impact on the genetic structure of the population.

Secondly, when the metapopulation is at stationary state, the asymptotic allele frequency *p*^*^ does not depend on the fecundity parameters *f*_0_ and *β*. However, the allele frequency crucially depends on the maturation rate *m* as well as the survival probabilities *s_J_* and *s_A_*. In particular, increasing the maturation time which corresponds to reducing *m*, tends to increase the asymptotic frequency of alleles, which have initially a higher frequency among juveniles than among adults (*p_J_*(0) > *p_A_*(0)), otherwise it decreases its frequency (*p_J_*(0) < *p_A_*(0)).

#### Allele dynamics in a metapopulation at periodic steady states

Our result also applies when the metapopulation has reached a periodic steady state. This situation may occur when the intrinsic fecundity *f*_0*k*_ is large (see Fig. 1). In this situation, the the allele frequency among stages in each population will stabilize around the constant value *p*^*^ which does not depend on time. However, this quantity crucially depends on the initial configuration of the metapopulation **N**\*(0). Since **N**\*(*t*) is T-periodic, the initial value **N**\*(0) can take *T* different values. Thus in a time varying metapopulation, the long time behaviour of an allele **n** crucially depends on when this allele appears in the metapopulation (see Appendix B.2 and Fig. 9 for more details).

Moreover, under this dynamics of the metapopulation, our numerical simulations (see Fig. 9) show that the asymptotic allele frequency *p*^*^ does depend on the fecundity of the species. As a result, we can conclude that the fecundity only plays a role when the metapopulation densities vary in time.

#### Comparison with a stochastic individual–based model

We compare our analytical results with the outcome of an individual-based model (IBM model) in large population described in appendix C. In the IBM model, each allele reproduces, survives and matures with the probability given by the reproduction matrix **F**_*k*_ defined by (7) and disperses with probabilities given by the dispersal matrices **D**_*kl*_ defined by (6). In our IBM model, the size of each population composing our metapopulation is finite of order *N_s_β_i_* where *N_s_* is a parameter of the IBM model describing the typical size of the population. When the population sizes are large (*N_s_β_j_* ≈ 5.10^3^), our deterministic model provides a good approximation of the stochastic model in various situation: metapopulation that either stabilizes around an equilibrium or a periodic steady state (see Fig. 3). In this regime of large population size, the genetic drift only occurs at really large time scale (see Fig. 11 and Fig. 12 in Appendix C for more details).

### 3.2 Allele dynamics in a non equilibrium metapopulation: linear case

We assume here that the metapopulation is no longer at steady state initially in order to capture the effect of transient dynamics of the metapopulation on the allele dynamics. However, in this

section, we do not assume any density–dependence in the metapopulation dynamics, that is thereproduction matrices **F_*k*_** are constant and do not depend on the population size **N_*k*_** in the habitatk. In particular, the hypothesis (*H*_2_) may be not satisfied. In this situation, the model (1) becomes linear and we know from classical metapopulation theory, that the proportion of each stage in each habitat of the metapopulation will converge to an asymptotic proportion. Moreover, from the linearity of the model, the metapopulation **N** and any allele **n** in this metapopulation satisfy the following linear model:

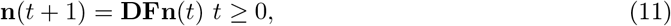

where **D** and **F** satisfy hypothesis (*H*_1_), (*H*_3_) and (*H*_4_) and initially we have 0 ≤ *n_j,k_*(0) ≤ *N_j,k_*(0) for all (*j*,*k*) ∈ {1,…,*ω_s_*} × {1,…,*ω_h_*}.

Using the properties of the dispersal and the reproduction matrix, we have the following result:

#### Theorem 2.

*Let* **N** *and* **n** *be solutions of* (11) *starting from a non-negative initial condition* **N**(0) *and* **n**(0) *such that* 0 ≤ *n_j,k_*(0) ≤ *N_j,k_*(0) *for all* (*j*, *k*) ∈ {1,…, *ω_s_*} × {1,…,*ω_h_*}. *Then*

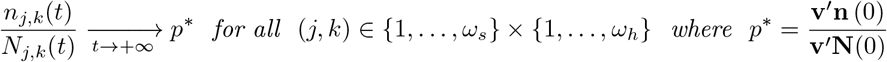

*and* **N**\* *and* **v** *are respectively the eigenvectors of the matrix* **DF** *and of its transpose associated to the principal eigenvalue* λ > 1.

Our result, proved in section 7.2, shows that any allele frequency converges to an asymptotic positive frequency thanks to the primitivity property of the projection matrix **DF** even if the metapopulation is not initially at equilibrium. Moreover, the convergence occurs geometrically fast with rate given by the ratio between the principal eigenvalue and the maximum of absolute value of the remaining eigenvalues of **DF**. We can also observe that the asymptotic allele frequency *p*^*^ crucially depends on the initial repartition of the metapopulation **N**(0) as in the periodic case.

## 4 Effect of life–history traits on neutral genetic diversity

Our previous results have provided some insights on the genetic richness of a metapopulation which may either be at equilibrium or may vary periodically in time. Now, we aim to understand the effect of the life-history traits (juvenile and adult survival probabilities, maturation probability and fecundity) on the diversity both at local habitat scale through the *α*–diversities among habitat and among stage and at global scale through the *γ*–diversity. Our previous results show that asymptotically in time, these diversity indices at both local and global scales are identical. Thus, if the initial metapopulation is composed of *I* alleles with density **n**^*i*^, the diversity indices *γ-Div*(*t*), (*α-Div*)_*h*_(*t*) and (*α-Div*)_*c*_(*t*) defined respectively by (3), (4) and (5), will converge towards the following asymptotic diversity index Div:

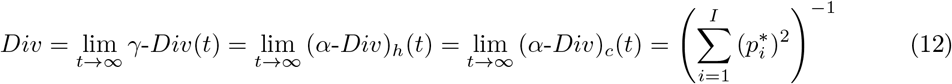

where 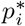 are the asymptotic allele frequencies given by Theorem 1 associated with initial allele densities **n**^*i*^(0).

In this section, we focus on the effect of life-history traits on the genetic diversity in a single population. The general case of a metapopulation will be investigated in the following section. Here, we look at most basic life cycle division between juveniles and adults. The reproduction matrix is thus described by (7) and it is characterized by 4 traits: the fecundity *f*, the maturation probability m and the juvenile and adult survival probabilities *s_J_* and *s_A_*.

This simple model permits us to examine four classes of life-histories, depending on the reproduction strategy and the development rate. The reproduction strategy ranges from semelparous (reproducing once in lifetime) to iteroparous (reproducing repeatedly). Semelparity is obtained when *s_A_* → 0 and iteroparity when *s_A_* > 0. We can also distinguished different types of development from precocious (rapid development to maturity, *m* → 1) to delayed (*m* < 1). The combination of these two dichotomies provides four classes of life-histories:

- Precocious semelparity: e. g., many annual plants and insects with rapid development and only one reproductive event.
- Precocious iteroparity: e. g., small mammals and birds, which begin reproducing when a year or less old, but may survive and reproduce for several years.
- Delayed semelparity: e. g., periodical cicadas or periodically-flowering bamboos that live for many years before maturity, and then reproduce only once.
- Delayed iteroparity: e. g., humans, whales, other large mammals, and some birds (albatrosses) that have long pre-reproductive periods and then survive and reproduce for many years.

Using our previous results, we aim to understand how the neutral genetic diversity is shaped by the following life-history traits: (1) the development duration which depends on the maturation probability *m*, (2) the survival probabilities *s_A_* and *s_J_* and (3) the fecundity f. To quantify the effect of those traits on the neutral genetic diversity, we assume that the population is at stationary equilibrium and it is composed of two alleles with initially different frequencies among each population stage. More precisely, we assume that initially, one allele is carried only by juveniles while the other one is carried by adults, ie the first allele have a frequency of *p_J_*(0) = 1 among juveniles and *p_A_*(0) = 0 among adults. Such situation may occur, for instance, if we take two monomorphic populations at equilibrium with different alleles in each population and we exchange the adults between the populations. Thus we obtain two populations at equilibrium with two different alleles in each population and initially each allele is carried only by one stage among the populations. Thus at the beginning, the diversity among each stage is lower than the diversity in the population. Since individuals maturate, reproduce and eventually die, in each population, the allele densities will evolve over time according to the model (2). Moreover, this situation allows us to investigate the effect of the different stages on the diversity.

### 4.1 Does long juvenile stage promote neutral genetic diversity?

We first investigate the effect of the juvenile stage on the diversity. More precisely, we focus on the maturation probability *m* which influences the duration of the juvenile stage. Indeed, we know from our model that the mean duration of the juvenile stage is *T_J_* = 1/(1 – (1 – *m*)*s_J_*)). When the maturation probability is small (delayed development *m* < 1), the juvenile stage lasts for several generations, while if *m* is large (precocious development *m* → 1), the juvenile lifetime reduces.

In this section, we only focus on population at equilibrium, that is **N**\* is constant over time. We know from our previous results that the asymptotic diversity Div is given by the following analytical expression

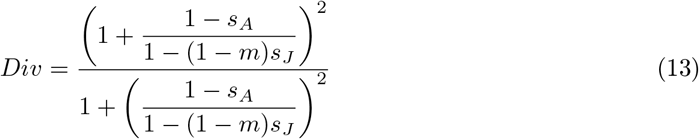

Then we get the following properties on the diversity.

#### Proposition 1.

*Let* **N**\* *be the equilibrium solution of* (1) *composed of two alleles* **n**^1^(0) = (*J*\*, 0) *and* **n**^2^(0) = (0, *A*\*). *Then the asymptotic diversity* Div *defined by* (12) *satisfies the following properties*:

- *if s_J_* < *s_A_ then Div is decreasing with respect to the maturation probability m*;
- *if s_J_* ≥ *s_A_ then Div attains a maximum at* 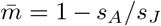 *and* Div *is increasing with respect of maturation m if* 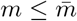 *and Div is decreasing with respect of maturation m if* 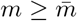.

This result shows that the effect of maturation time on diversity truly depends on the reproduction strategy of the species (semelparous or iteroparous) through the adult survival probability. Among iteroparous species, diversity is enhanced when the juvenile lifetime is long (m < 1). Thus, we should find higher diversity among delayed iteroparous species than among precocious iteroparous species (see blue curve in Fig. 4(a)). This beneficial effect of the juvenile stage on genetic diversity has already been observed in plants (Austerlitz et al., 2000).

**Figure 4:**
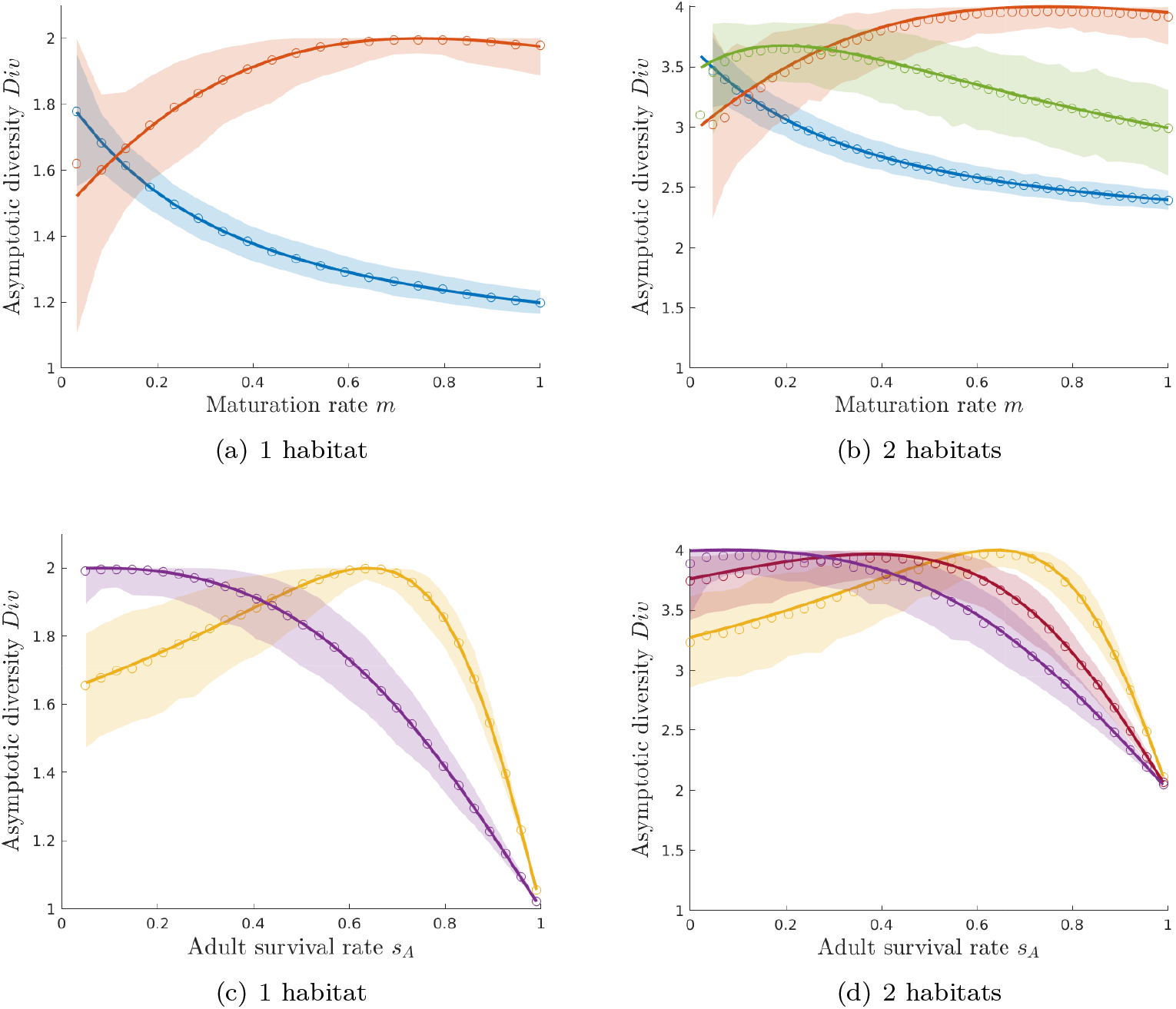
Effect of maturation probability *m* and adult survival *s_A_* on asymptotic diversity *Div*: (a)-(c) in a single population and (b)-(d) in metapopulation living in two habitats (migration rates are *ε*_12_ = *ε*_21_ = 0.1) for the deterministic model (plain curves) and the IBM model for *N* = 50 individuals (circle marked curves correspond to the medians and the shaded regions correspond to the 99% confidence intervals over 10^3^ replicates). Each colour corresponds to different sets of parameters: blue curves *s_A_* = 0.2, red curves *s_A_* = 0.9 and green curve *s*_*A*1_ = 0.2 and *s*_*A*2_ = 0.9; orange curves *m* = 0.2, purple curves *m* = 0.9 and cyan curve *m_1_* = 0.2 and *m*_2_ = 0.9. Habitat characteristics: *s*_*J*1_ = *s*_*J*2_ = 0.8, *f*_01_ = *f*_02_ = 10 and *β*_1_ = *β*_2_ = 150.

Conversely, among semelparous species, a shorter juvenile lifetime (*m* → 1) will promote diversity (see red curve in Fig. 4(a)). More precisely, we can observe that the diversity depends on the ratio between the juvenile lifetime *T_J_* and the adult lifetime *T_A_* = 1/(1 – *s_A_*). In particular, the diversity is maximal when the two lifetimes are similar. Conversely, when the lifetimes are very different, the diversity erodes. Indeed, when the adults’ lifetime is longer than the juveniles’ lifetime, the adults last for a longer time and thus they produce many individuals with similar alleles, which unbalanced the allele frequencies and erodes diversity. Similarly, when adult lifetime is shorter than juvenile lifetime, the replacement rate of juveniles by reproduction is low, which also reduces the diversity. Thus we show that the diversity does not really depend on the short or long lifespan of a species rather than the lifetime of its different stages along its life.

### 4.2 Does adult survival promote neutral genetic diversity?

Using the previous formula and Proposition 1, we now investigate the impact of adult survival on the neutral genetic diversity. In a population at equilibrium, we show that diversity is negatively dependent on the adult survival. In particular, immortal species should have a small diversity (*Div* → 1 when *s_A_* → 1 see Figure 4(c)). And the diversity reaches a maximum when adult survival satisfies 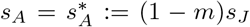. We observe as before that the maximum of diversity is reached when the lifetime of juvenile and adult are identical. In addition, we see that an increase of adult survival among species with precocious development has a detrimental effect on its diversity while species with delayed development need a large adult survival. Thus, precocious semelparous species have higher diversity than precocious iteroparous species (see red curve Figure 4(c)). While delayed semelparous species have lower diversity than delayed iteroparous species (see blue curve Figure 4(c)).

### 4.3 Fecundity only affects diversity in time varying populations

We now investigate the effect of fecundity on the neutral genetic diversity of a population either at equilibrium or periodically varying in time. First, we can see from formula (13) that the diversity of a population at equilibrium does not depend on the fecundity. This unexpected result was already observed experimentally among animals (De Kort et al., 2021).

However, when the population size varies periodically in time, which may result from a high fecundity, the diversity does depend on the fecundity (see Fig. 5). Moreover, the diversity might have different values depending on the initial configuration of the population (see dots in Fig. 5). As a result, we show that on average, the increase of fecundity drastically reduces the diversity among semelparous species while it has no significant effects on iteroparous species (see dashed curves in Fig. 5).

**Figure 5:**
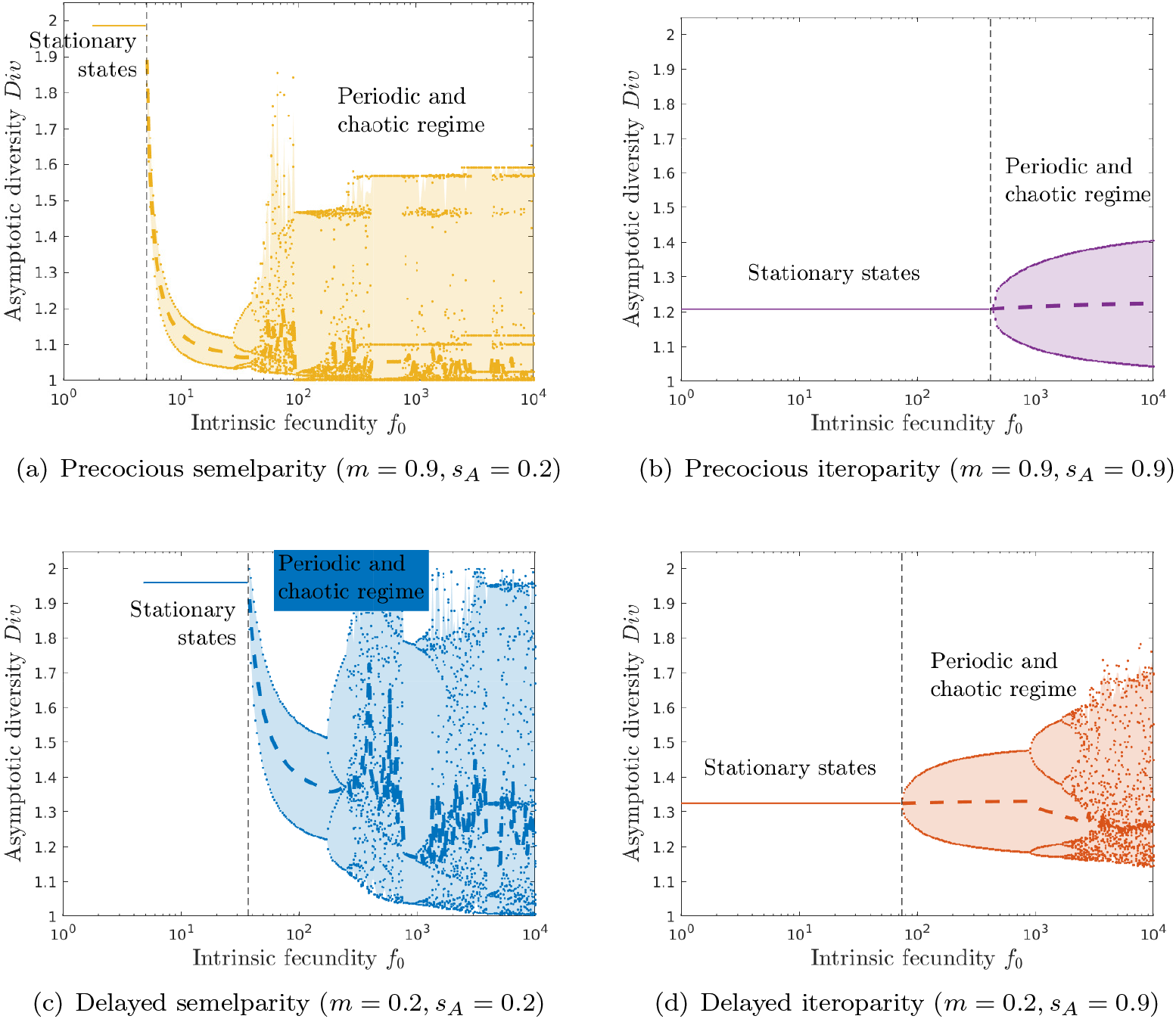
Effect of fecundity *f*_0_ on asymptotic diversity Div in a single population for the deterministic model. Each colour corresponds to the four different classes of life-histories: (a) precocious semelparity (orange *m* =0.9, *s_A_* = 0.2), (b) precocious iteroparity (purple (*m* = 0.9, *s_A_* = 0.9)), (c) delayed semelparity (blue (*m* = 0.2, *s_A_* = 0.2)) and (d) delayed iteroparity (red (*m* = 0.2, *s_A_* = 0.9)). Straight lines correspond to the equilibrium regime, dots correspond to the different diversity values in the periodical and chaotic regimes and dashed curves correspond to the mean values of diversity under those regimes. Habitat characteristics: *s*_*J*1_ = 0.5 and *β* = 150.

## 5 Intertwined effect of dispersal and life-history on diversity

An other important process which structures the genetic diversity in a metapopulation is the dispersal. In this section, we focus on a metapopulation composed of two habitats with possibly different characteristics corresponding to different life-histories, connected through dispersal. We show that dispersal can mitigate effect of the life-histories at the metapopulation scale. In addition, we show that difference in migration among stages can significantly modify the diversity.

### 5.1 Dispersal can modify the effects of life-histories

In a single population, we have shown analytically that the longer juvenile stage promotes diversity if adult survival is large while it reduces diversity when adults survival is low (see Proposition 1). Similarly, we have shown analytically that a large adult survival promotes diversity among species with delayed development (small maturation rate), while it reduces drastically the diversity with precocious development (large maturation rate). The same patterns occur when the habitats are identical (see red and blue curves in Fig. 4(b) and orange and purple curves in Fig. 4(d)).

However, when habitat characteristics are different, the migration between habitats balances the antagonist effects of the juvenile stage and the adult survival. Moreover, we can see that the dispersal may even enhance the diversity when the habitats are heterogeneous. The synergy occurs when juvenile stage is long or when adult survival is intermediate (see green and dark red curves above the others in Fig. 4(b)–4(d)).

### 5.2 Does migration always promote diversity?

We now investigate the effect of dispersal in a metapopulation living in two habitats with different characteristics. More precisely, they differ either in maturation probability *m* or in adult survival *s_A_*.

When the migration between the habitats is symmetric (*ε*_12_ = *ε*_21_), then it generally promotes diversity (see Fig. 6(a)). In particular, when populations have different maturation rates between habitats, the dispersal has no effect on iteroparous species (see blue curve in Fig. 6(a)) while it reduces diversity if migration is too high among semelparous species (see red curve in Fig. 6(a)). However, when the adult survival probability is different depending on the habitat, the dispersal always enhances diversity (see orange and purple curves in Fig. 6(a)).

**Figure 6:**
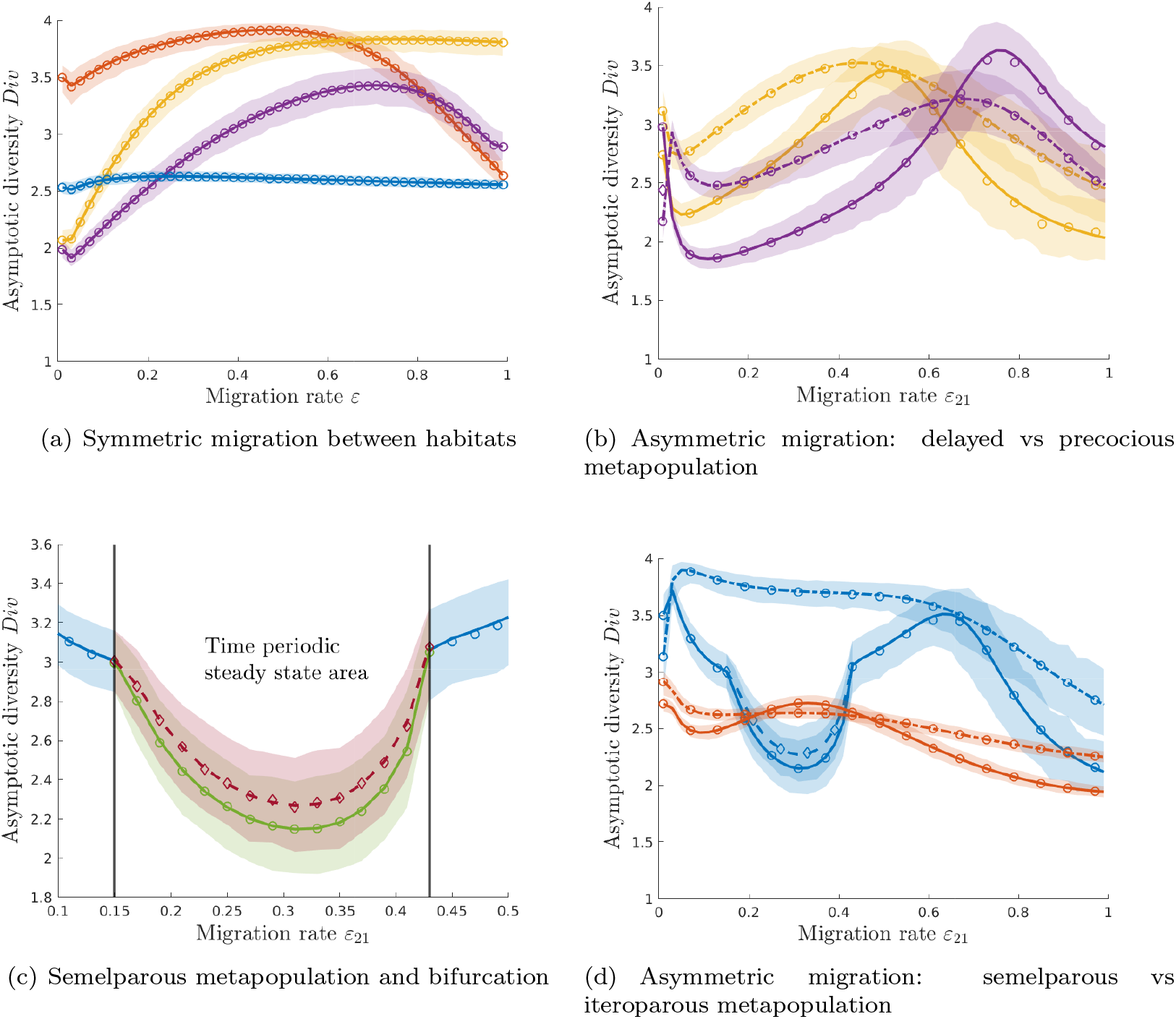
Effect of migration rates *ε_kl_* on asymptotic diversity *Div* for the deterministic model (plain curves) and the IBM model for *N* = 50 individuals (circle marked curves correspond to the medians and the shaded regions correspond to the 99% confidence intervals over 10^3^ replicates). Each colour corresponds to different set of life-history parameters: blue curves semelparous species (*s*_*A*1_ = *s*_*A*2_ = 0.9, *m*_1_ = 0.2 and *m*_2_ = 0.9), red curves iteroparous species (*s*_*A*1_ = *s*_*A*2_ = 0.2, *m*_1_ = 0.2 and *m*_2_ = 0.9), orange curves delayed development (*m*_1_ = *m*_2_ = 0.2, *s*_*A*1_ = 0.2 and *s*_*A*2_ = 0.9) and purple curves precocious development (*m*_1_ = *m*_2_ = 0.9, *s*_*A*1_ = 0.2 and *s*_*A*2_ = 0.9). When migration is asymmetric (b)-(d): plain curves correspond to low migration (*ε*_12_ = 0.05) and dashed dotted curves to high migration (*ε*_12_ = 0.2). In area where the steady state is time periodic with period *T* = 2 (c), each colour corresponds to the diversity associated to one of the two values of the steady state. Habitat characteristics: *s*_*J*1_ = *s*_*J*2_ = 0.8, *f*_01_ = *f*_02_ = 1.5 and *β*_1_ = *β*_2_ = 150.

Now, we focus on asymmetric migration between heterogeneous habitats. When adult survivals are different, a high migration probability from the habitat with larger adult survival will promote diversity for any duration of juvenile stage (see Fig. 6(b)).

When the maturation rates are different, the effect of the migration depends on the species reproduction trait. If the species is iteroparous, migration has small effect on diversity (see red curve in Fig. 6(d)). However, the diversity is higher when individuals are more likely to move to habitat with low maturation probability. Thus for iteroparous species, individuals should remain in habitat with longer juvenile stage.

Conversely, among semelparous metapopulation, individuals should move to habitat with a high maturation probability (see blue curves in Fig. 6(d)). In addition, when the migration rates are low, the equilibrium dynamics of the metapopulation changes from stationary equilibrium to periodically varying steady state. Under the time varying scenario, diversity decreases drastically and multiple values can be achieved (see blue curves in Fig. 6(c)).

### 5.3 Effect of individuals migration on diversity

We now look at the effect of the migration of each stage in the metapopulation, that is 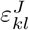 and 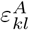 in the definition of the dispersal matrix. We assume that the migration is symmetric between habitats in the sense that 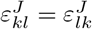 and 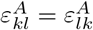 for all habitats *k*, *l* in {1, 2}. We investigate how the diversity responds to a change on the ratio of migration *ε^A^*/*ε^J^*. For each value of the migration ratio, we pick different values for the migration rates. Thus the mean migration probability between habitat is not constant for a given value of the migration ratio. However, we show that an heterogeneous migration probability between the stages of the metapopulation has critical impact on the diversity. In particular, we show that the iteroparous species need a higher migration probability from the adults than the juvenile to keep a higher diversity (see blue curve in Fig.7(a)). Conversely, among semelparous species, the migration rate of the juvenile needs to be higher than the migration rate of the adult to keep a high diversity (see red curve in Fig.7(a)). When the duration of the juvenile stage is long which corresponds to a delayed development, the diversity is higher when the adults disperse more than the juveniles (see orange curve Fig.7(b)). When the juvenile stage duration is reduced, the diversity is high when only the adults disperse or when only the juveniles disperse (see purple curve Fig.7(b)). In this case, two antagonist strategies emerge to keep a high diversity. In addition, we can observe that the diversity is more variable when the adults disperse more than juveniles.

**Figure 7:**
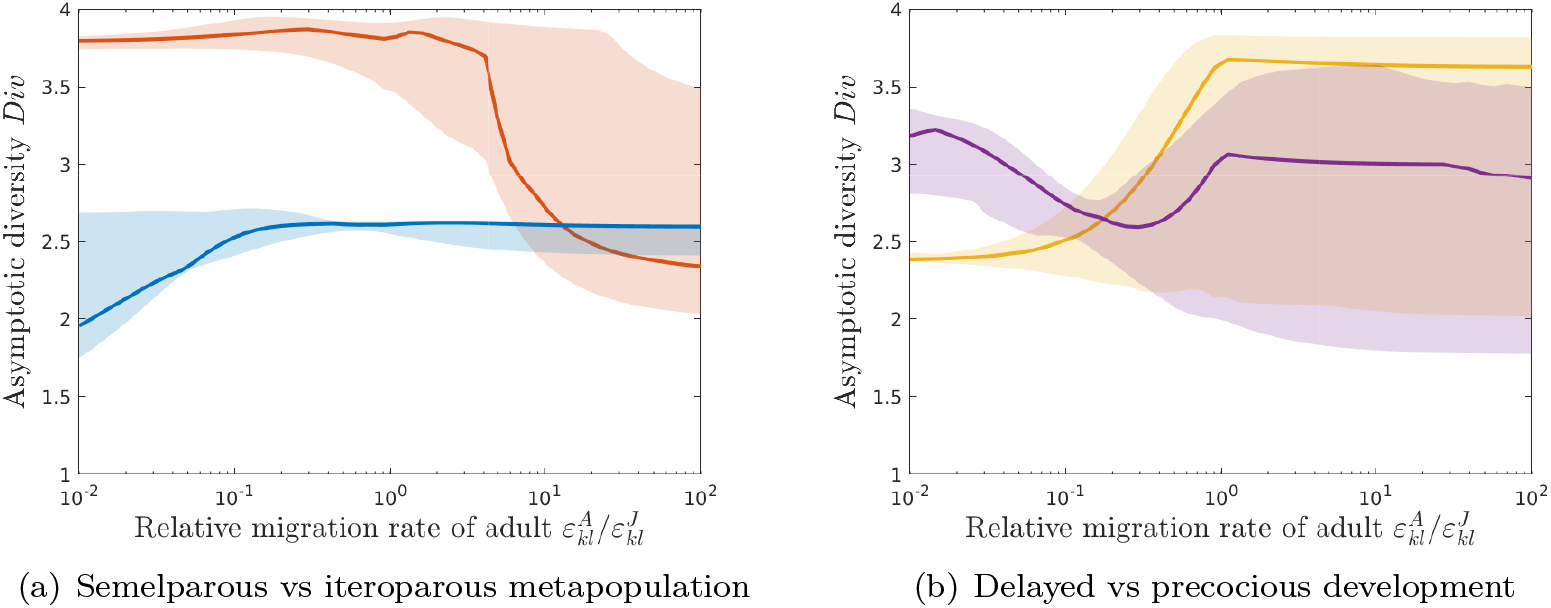
Effect of the migration ratio between stages 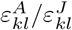 on the asymptotic diversity *Div* for four life-history parameter sets defined in Fig. 6. Plain curves corresponds to the median of the asymptotic diversity *Div* over 100 couples of migration rate 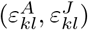 whose constant ratio ranges from 10^-2^ to 10^2^.

**Figure 8:**
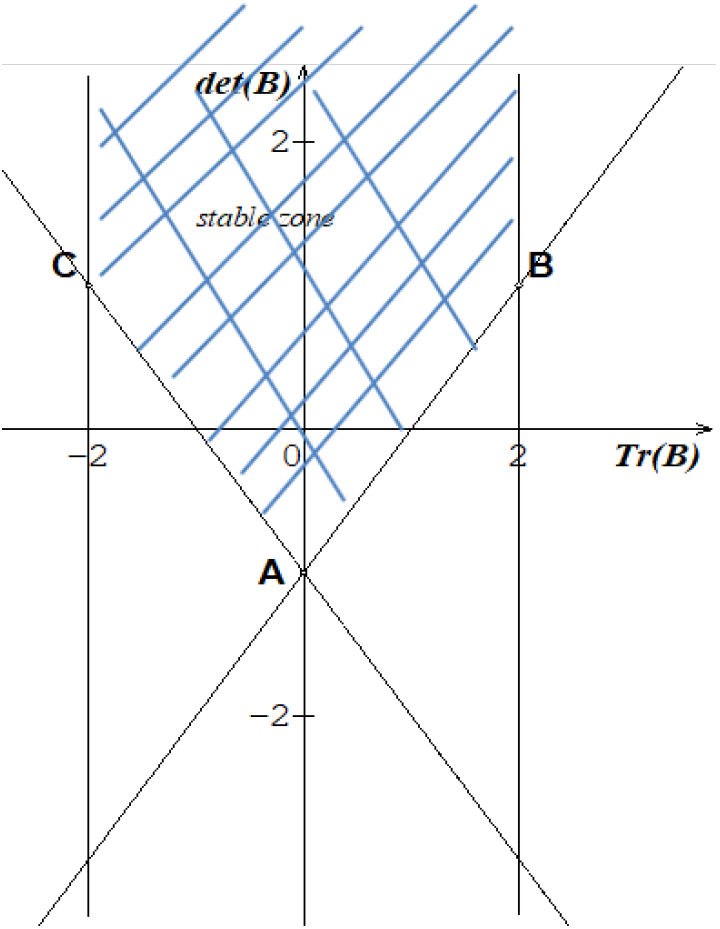
the stable zone corresponds to the hatched area

**Figure 9:**
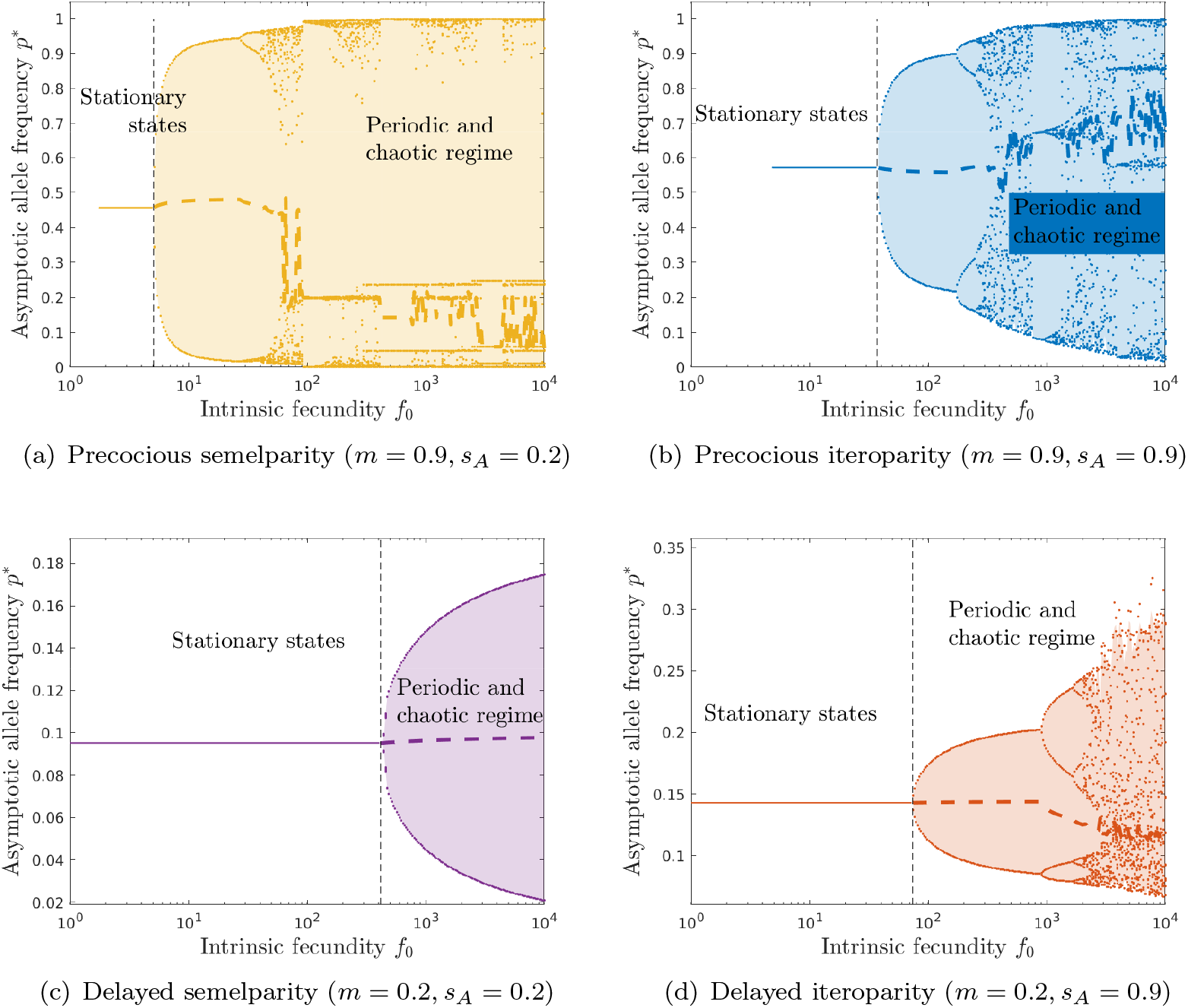
Effect of fecundity *f*_0_ on asymptotic diversity *Div* in a single population for the deterministic model. Each colour corresponds to the four different classes of life-histories: (a) precocious semelparity (orange *m* = 0.9, *s_A_* = 0.2), (b) precocious iteroparity (purple (*m* = 0.9, *s_A_* = 0.9)), (c) delayed semelparity (blue (*m* = 0.2, *s_A_* = 0.2)) and (d) delayed iteroparity (red (*m* = 0.2, *s_A_* = 0.9)). Straight lines corresponds to the equilibrium regime, dots corresponds to the different diversity values in the periodical and choatic regimes and dashed curves corresponds to the mean values of diversity under those regimes. Habitat characteristics: *s*_*j*1_ = 0.5 and *β* = 150.

**Figure 10:**
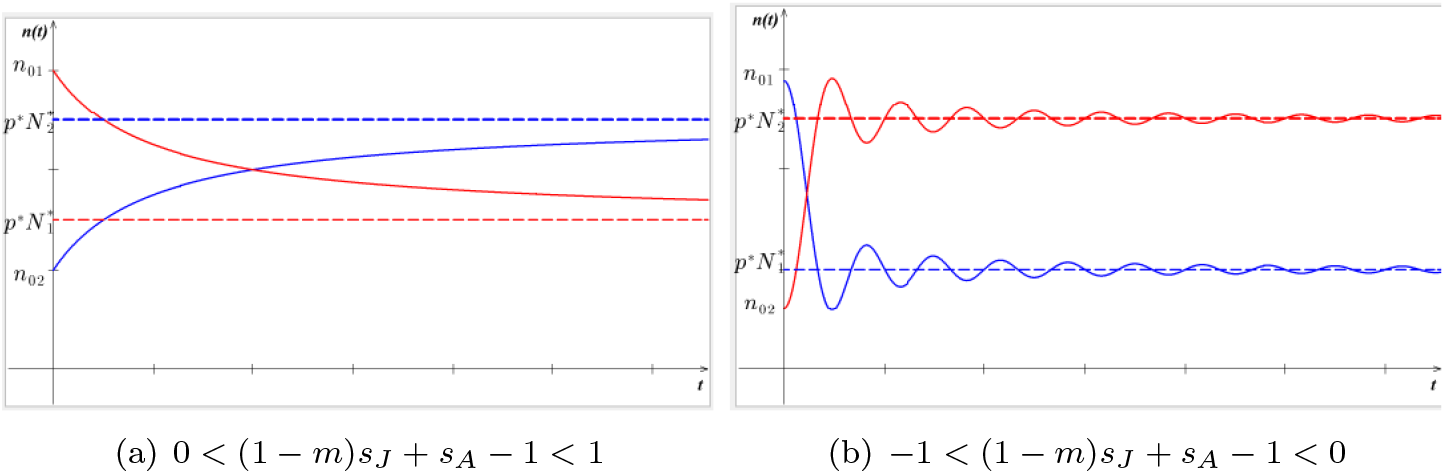
Dynamics of the allele **n**(*t*) in a single habitat with either: (a) monotonic convergence toward the stationary state or (b) convergence with damped oscillation around the stationary state.

**Figure 11:**
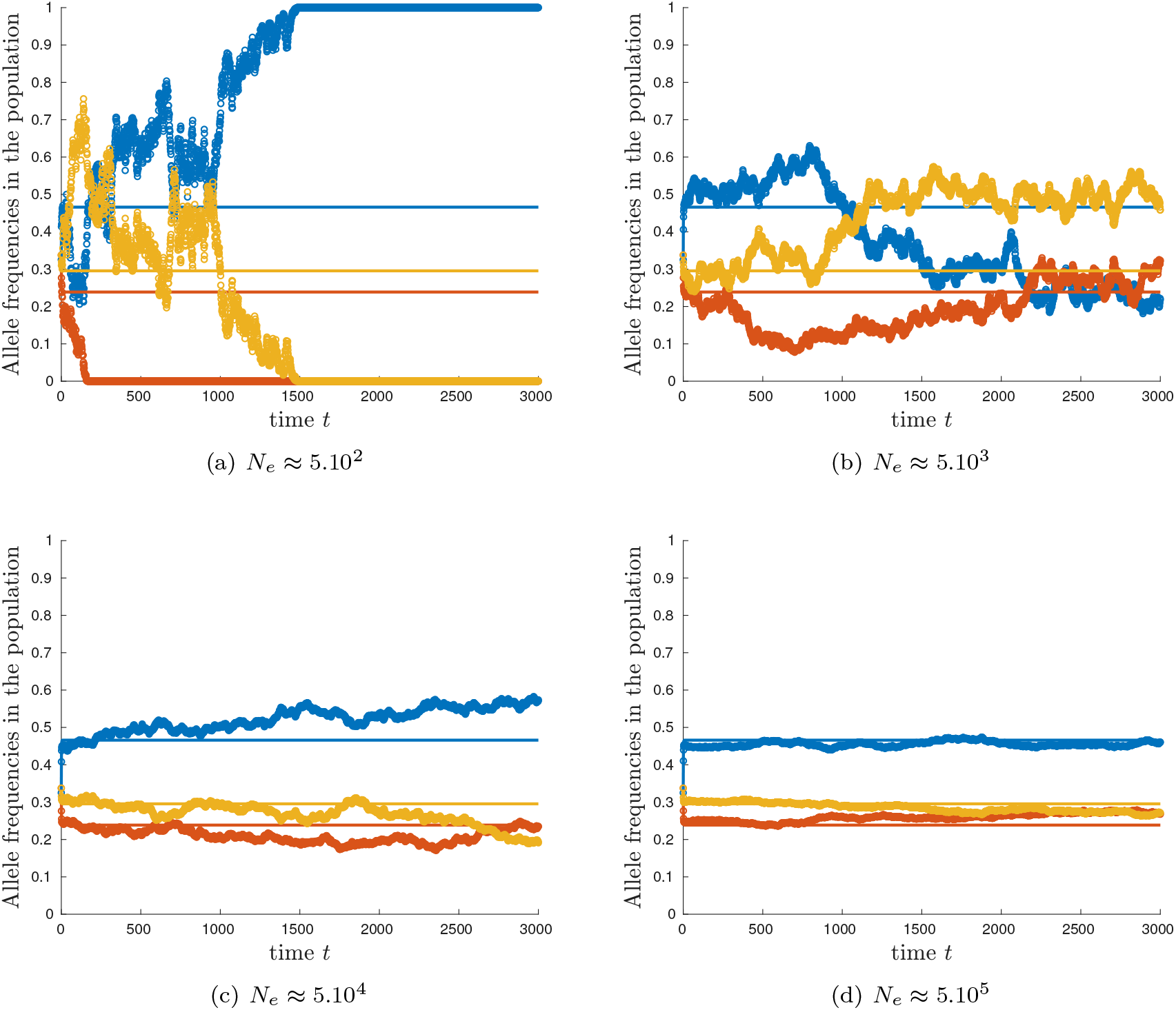
Temporal dynamics of the allele frequency in the population of 3 alleles in a single habitat for the deterministic model (plain lines) and the IBM model with (a) 5.10^2^, (b) 5.10^3^, (c) 5.10^4^and (d) 5.10^5^ individuals (circle marked curves correspond to one trajectory of the IBM model). Habitat characteristics: *f*_01_ = 1.5, *s*_*A*_1__ = 0.7, *s*_*J*_1__ = 0.8, *m*_1_ = 0.2, *β*_1_ = 100 and *N_s_* = 50.

**Figure 12:**
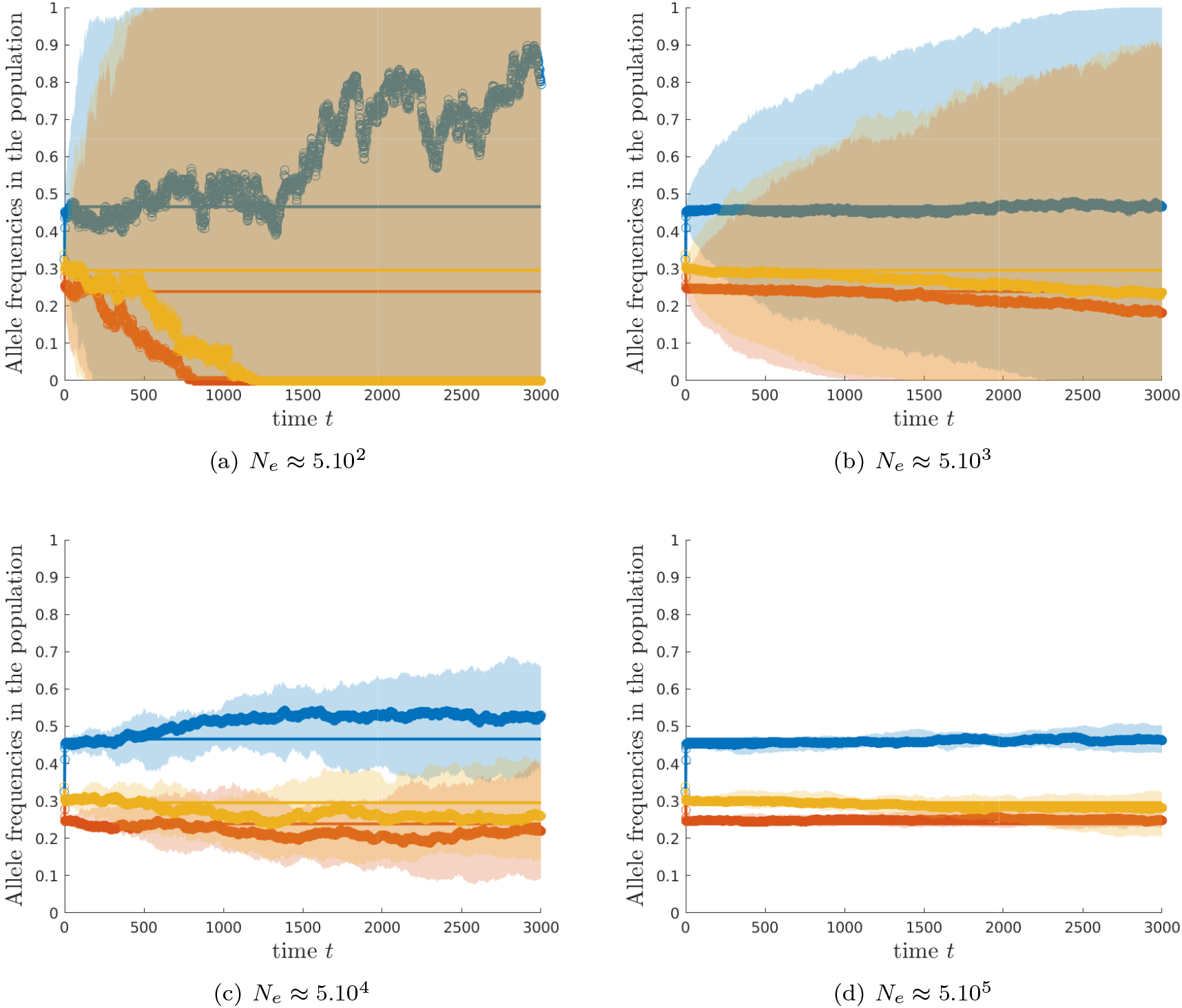
Temporal dynamics of the allele frequency in the population of 3 alleles in a single habitat average over 10^3^ replicates. Plain curves correspond to the deterministic model. Circle marked curves correspond to the medians and the shaded regions correspond to the 99% confidence intervals over 10^3^ replicates of the IBM model with: (a) 5.10^2^, (b) 5.10^3^, (c) 5.10^4^ and (d) 5.10^5^ individuals. Habitat characteristics: *f*_01_ = 1.5, *s*_*A*_1__ = 0.7, *s*_*J*_1__ = 0.8, *m*_1_ = 0.2, *β*_1_ = 100 and *N_s_* = 50.

## 6 Conclusion and discussion

In the present work, we investigate the effect of dispersal and life-history traits of different populations composing a metapopulation on its neutral genetic diversity at different spacial scales using a deterministic mathematical model. Our model is a classical metapopulation model (Holt,1985) combined with a matrix projection model in discrete time which takes into account the life-history traits and the stage–structure of the populations (Neubert and Caswell, 2000). Extending the inside dynamics approach developed by Hallatschek and Nelson (2008), Garnier et al. (2012) and Roques et al. (2012), we describe the dynamics of each neutral allele composing the metapopulation. We both consider a metapopulation that has already reached an equilibrium or a metapopulation that evolves in time. In particular, for the linear model, we deal with a metapopulation that is not initially at equilibrium and we describes the transient as well as the long time behaviour of the metapopulation and its neutral genetic diversity. In addition, for the nonlinear model, we are able to describe the entire dynamics of neutral genetic alleles in a metapopulation at stationary

equilibrium as well as in a metapopulation that fluctuates periodically in time. Our analytic characterization of the proportion of each neutral allele for large time provides us quantitative insights on the effect of life-history traits and dispersal on the neutral genetic diversity of the metapopulation at local and global scale. Moreover, our deterministic approach agreed with the dynamics of classical individual–based models.

First we have shown that the stage-structure of the populations influences the asymptotic proportion of each allele even in a single habitat. In particular, if several alleles are equally distributed in the population but with different proportions in each stage then asymptotically, their proportions will be different. Our results agree with the experimental results obtained on the freshwater herbivore *Daphnia pulex* (Nelson et al., 2005).

We also show that some life history traits truly influence the diversity. However, their effect depends on both the reproductive strategy (semelparous or iteroparous) and the development strategy (precocious or delayed). In particular, the presence of juvenile stage in expanding population is known to have profound impact on genetic diversity (Austerlitz et al., 2000; Bonnefon et al., 2013; Marculis et al., 2019). With our mathematical approach, we are able to quantify the impact of each demographic parameter on the diversity of a single population. We show that a long juvenile stage promotes diversity among iteroparous species. This beneficial effect of a long juvenile stage was already observed among plants (Austerlitz et al., 2000), birds (Eo et al., 2011) or mammalians (Doyle et al., 2015). Among those long-lived species, the long juvenile stage slows down the reproduction so that all individuals can contribute to the diversity of the population. Conversely, among semelparous species, precocious development will promote diversity. This beneficial effect of maturation was observed among fishes (Williams, 1985; Valiente et al., 2005; Dalongeville et al.,2016). Semelparous precocious species reproduce earlier and have larger number of descendant leading to rapid genetic mix. From our mathematical analysis, we show that this antagonist effect of the maturation results from the trade off between the lifetime of juveniles and adults. We show that diversity is optimal when the lifetime of juveniles and adults are similar. Thus, if adults last only for few generations (semelparous species), juveniles should maturate quickly (precocious development). Conversely, if adults are long lived (iteroparous species), the lifetime of juveniles should be long. Thus although species longevity significantly influences animals and plants diversity (De Kortet al., 2021; Romiguier et al., 2014), it cannot explain the diversity by itself.

We also show that the survival rate of adult have significant effect on diversity. In particular, among species with a delayed development or maturation, a degradation of the environment, characterized by a decrease of survival probabilities, will erode diversity. Actually, among mammalians, diversity is known to be lower in threatened and endangered species than in least concern species (Doyle et al., 2015). Conversely, species with precocious development will maintain higher diversity under harmful conditions. It has already been observed experimentally on wild salmon (Valiente et al.,2005). They show that diversity does not depend on latitude. However, maturation increases with latitude and environment becomes less favourable with small latitude. It agrees with our result showing that maturation rate can counterbalance diversity loss when adult survival rate decreases. As result, we show that reproductive strategy may well explain genetic diversity.

Fecundity is an other important life history trait which may significantly impact the genetic diversity. In particular, among time fluctuating semelparous populations, we show that an increase of fecundity dramatically erodes diversity. However, among time fluctuating iteroparous populations, its effect is less significant and populations with high fecundity may harbour higher genetic diversity. Furthermore, among stable population at equilibrium, we show that fecundity has no significant influence on genetic diversity. Recently,De Kort et al. (2021) also show that fecundity have a significant effect on animals genetic diversity only in endemic species while it has no significant effect in general. Thus the effect of fecundity depends on the biogeography as well as the population dynamics.

We also recover that the dispersal behaviour has profound impact on neutral genetic structure of a metapopulation (Wright, 1949; Lynch, 1988). We first show that dispersal between populations with different life-history traits may balance the antagonist effect of those traits and thus promotes genetic diversity. In particular, dispersal enhances diversity between populations with delayed maturation or between short-lived iteroparous populations. This beneficial effect of dispersal has also been observed among plants’ species range. Dispersal can moderate the decline in population genetic diversity from the core to edge habitats predicted from the biogeography theory (De Kort et al., 2021).

However, we show that increasing or decreasing the dispersal over one habitat might have detrimental consequences on diversity. This unbalanced dispersal might occur when individuals try to escape from bad quality habitat in the context of environmental change (Jenouvrier et al., 2017). In particular, if the dispersal from a good habitat to a less favourable habitat is small while the dispersal in the opposite direction is really large then the diversity is low (Garnier and Lafontaine,2021). This situation shows that escaping from bad habitat to survive might endanger genetic diversity even if it improves the persistence of the metapopulation by enhancing its size (Holt, 1985).

However, this detrimental effect of asymmetric dispersal might be compensated by different dispersal behaviours between juveniles and adults. We show that the dispersal ability of the different stages also have an impact on the genetic diversity. In the situation of a good habitat and a bad habitat, a symmetric dispersal of the juveniles might compensate the diversity erosion due to the massive adult immigration from the bad habitat to the good habitat. Indeed, the symmetric dispersal will help the allele mixing in the two habitats, which increases diversity. Such dispersal behaviour might occur if adults can use personal and social information to decide whether to leave a natal or current breeding site and where to settle (e.g. (Doligez et al., 2002)). Such ‘informed dispersal’ strategies (Clobert et al., 2009) enable individuals to settle in habitats of better quality. However, since it requires life experiences, we might expect that juveniles are less likely to use this dispersal strategy and thus disperse randomly in their environment. More generally, the effect of a discrepancy in dispersal behaviour between stages depends on the demographic characteristics of the different habitats. In particular, we show that adult dispersal promotes diversity when juvenile mortality is larger than adults mortality, while it has no effect when juvenile mortality is smaller than adult mortality. When the habitat becomes heterogeneous the effect of dispersal might change, but this effect remains small compare to the effect of global dispersal or demographic characteristics.

## 7 Proofs of the results

### 7.1 Proof of Theorem 1

#### The equilibrium case

Let **F** and **D** satisfy hypotheses (*H*_1_)-(*H*_4_) and **N**\* be a stationary state of equation (1), that is **N**\* = **DF** [**N**\*]**N**\*. Let **n**(*t*) be solution of (10) starting from **n**(0) such that 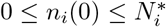 for all *i* ∈ {1,…, *m*} with *m* = *ω_c_* × *ω_h_*. Then

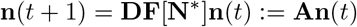

where the matrix **A** is primitive from hypothesis (*H*_4_).

Using the Perron-Frobenius theorem, we know that 1 is the principal eigenvalue of **A** with the positive eigenvector **N**\*. Let us denote (**p**_2_,…, **p**_*m*_) the following eigenvectors associated to the eigenvalues (λ_2_,…, λ_*m*_) such that |λ_*i*_| < 1 for all *i* ∈ {2,…, *m*} and the matrix **P** = (**N**\*, **p**_2_,…, **p**_*m*_). Then we can represent **n**(*t*) as follows:

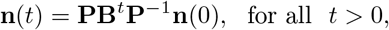

where **B** is a matrix of the following form with the eigenvalues on its diagonal:

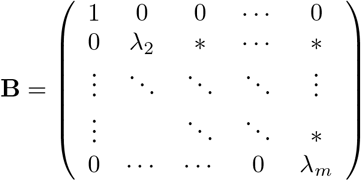

Then we deduce the following asymptotic behaviour of **n** as t → ∞:

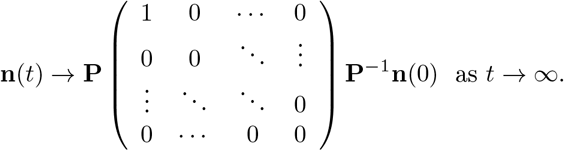

Moreover, we know that **P**^−1^ is the matrix where each row is equal to the eigenvectors (**v**, **v**_2_, …, **v**_*m*_) of the transpose matrix of **A**. Finally, we get

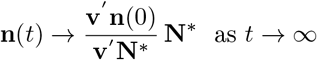

which concludes the first part of the theorem 1.

#### The periodic case

Let **F** and **D** satisfy hypotheses (*H*_1_)-(*H*_4_) and **N**\*(t) be a T-periodic steady state of equation (1), that is **N**\*(*t* + *T*) = **N**\*(*t*) for all *t* > 0. Let **n**(*t*) be solution of (10) starting from **n**(0) such that 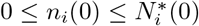 for all *i* ∈ {1,…, *m*} with *m* = *ω_c_* × *ω_h_*. Then

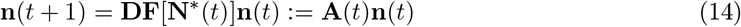

where the matrix **A**(*t*) is *T*-periodic since the steady state **N**\* is *T*-periodic. From the Floquet Theorem, we can decompose **n** as follows:

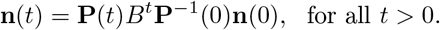

where **P**(*t*) is the *T*-periodic matrix constructed from **N**\* the T-periodic solution of (1) and (**p**_2_,…, **p**_*m*_) the *T*-periodic solutions associated to the Floquet exponents (λ_2_,…, λ_*m*_):

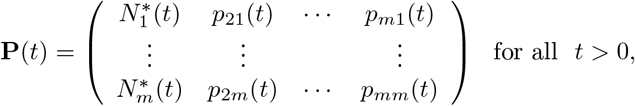

and the matrix **B** is of the following form with the Floquet exponents on its diagonal:

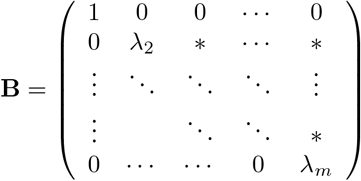

Moreover, we know that λ_*i*_ is a Floquet exponent if and only if 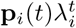 is a solution of (1), where **p**_*i*_ is a *T*–periodic solution, that is **p**_*i*_(*t* + *T*) = **p**_*i*_(*t*) for all *t* ≥ 0. Thus, we just need to find its first *T* component **p**_*i*_(0),…, **p**_*i*_(*T* – 1). These components satisfy the following eigenvalue problem:

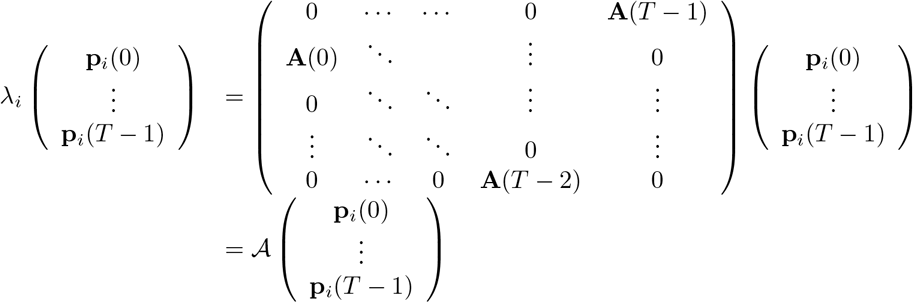

where 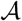 is a *T_m_* × *T_m_* matrix. Then, the eigenvalues of 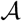 are the eigenvalues of **B** and the associated eigenvectors are the periodic fundamental solutions **P**(*t*). Thus we can characterize the periodic solutions **p** using this extended matrix 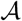.

The matrix 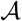 is non negative irreducible, then the Perron-Frobenius theorem insures that the eigenvalue 1 associated to the positive eigenvector **N**\* = (**N**\* (0),…, **N**\*(*T* – 1)) is the spectral radius of the matrix and it is simple. Moreover, its left eigenvector is also positive and simple. We also know that 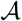 has *T* – 1 eigenvalues of absolute value 1 associated to eigenvectors which are circular permutations of **N**^*^ and all the other eigenvalues have absolute values strictly less than 1. We deduce that there is only one Floquet exponent of value 1 and all the other exponents are of absolute value less than 1, |λ_*i*_| < 1 for all *i* ∈ {2,…, *m*}.

We thus deduce the asymptotic behaviour of **n**:

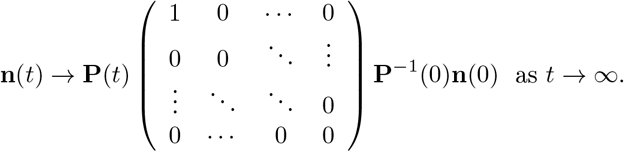

Then we can compute the matrix **P**^-1^(0) using the left eigenvectors of the matrix 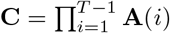. We can show that (**N**\*(0),**p**_2_(0),…,**p**_*m*_(0)) are the right eigenvectors associated to eigenvalues 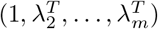 of the matrix **C**. From Perron-Frobenius theorem, all those eigenvectors are simple and there are left eigenvectors (**v**(0),**v**_2_(0),…,**v**_*m*_(0)) such that **v**(0) have all this components positive. Moreover, if we assume that (**v**(0), **N**\*(0)) = 1 and (**v***_i_*(0), **p**_*i*_(0)) = 1 for all *i* ∈{2,…, *m*}, then the matrix **p**(0) = (**v**(0)′, **v**_2_(0)′,…, **v**_*m*_(0)′)), where ′ is the transpose operator, satisfies **p**(0) = **P**^-1^(0). Then we eventually obtain that:

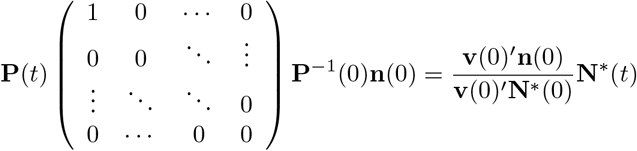

which concludes the proof of Theorem 1.

### 7.2 Proof of theorem 2, the linear case

We now consider the case where **F** does not depend on **N**. In this case, the metapopulation model becomes linear. Thus the metapopulation and any neutral allele **n** satisfy the following linear equation

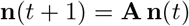

where **A** := **DF**, **D** and **F** satisfy hypothesis (*H*_1_)-(*H*_4_) and initially we have 0 ≤ *n_i_*(0) ≤ *N_i_*(0) for any *i* ∈ {1,…, *m*}.

Since **A** is a non negative primitive matrix, the Perron-Frobenius theorem provides the existence of a principal eigenvalue λ associated with a positive eigenvector **N**\*. Moreover, this eigenvalue is simple and the other eigenvalues λ_2_,…, λ_*m*_ satisfy |λ_*i*_| < λ for all *i* ∈ {2,…, *m*}. Moreover, we know that:

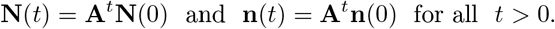

Then, using the eigenvector **v** of the transpose of the matrix **A** associated to the principal eigenvalue λ, we deduce from the Perron-Frobenius theorem that

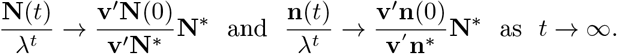

Then we conclude that each proportion of the neutral allele *n_i_*(*t*)/*N_i_*(*t*) satisfies the following asymptotic behaviour:

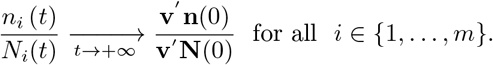

So, we see that the asymptotic proportion of this neutral allele in the metapopulation is given by:

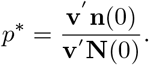

which concludes the proof of theorem 2.

### 7.3 The proof of proposition 1

Let us consider the equilibrium **N**\* in the case of only one habitat and two stages (juveniles *J* and adults *A*) with the special reproduction matrix **F** defined by (7). We have the following explicit expressions for **N**\* = (*J*\*, *A*\*):

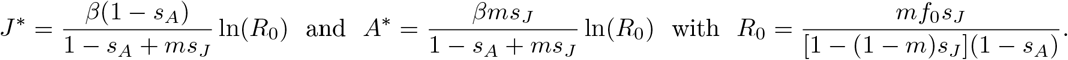

#### Behaviour of *p*\* with respect to the maturation rate *m*

Let us consider a neutral allele **n** starting with only juveniles, that is **n**(0) = (*J* *, 0). Our theorem 1 provides the following analytical expression for its asymptotic proportion *p*^*^

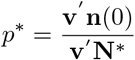

where **v** = (*v*_4_, *v*_2_) is the left eigenvector of **F** [**N**\*] associated to the eigenvalue 1. Using the explicit formula of **N**\*, we can compute analytically **F**[**N**\*] and **v**:

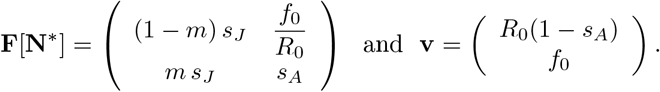

Then we have

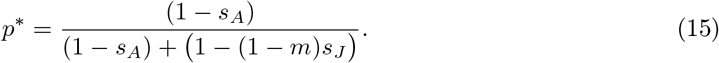

We can observe that the asymptotic proportion *p*^*^ is decreasing with respect to the maturation rate *m*.

#### Behaviour of the asymptotic diversity *Div* with respect to the maturation rate *m*

Now let us assume that the population is decomposed of two neutral alleles **n**_1_ and **n**_2_ starting respectively only with juveniles or adults, that is **n**_1_ = (*J*\*, 0) and **n**_2_ = (0, *A*\*). From the Theorem1and the previous computation, we have the analytical expression of the asymptotic proportions of each fraction, that is 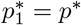, where *p*\* is defined by (15) and 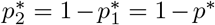. Thus we deduce the following asymptotic diversity *Div*

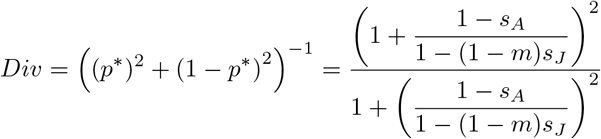

Thus, the derivative of *Div* with respect to the parameter *m* satisfies

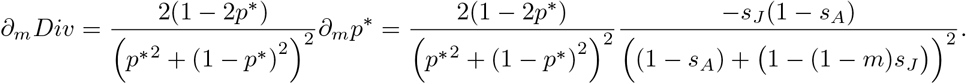

The sign of *∂_m_ Div* only depends of the sign of (2*p*\* – 1).

Using the explicit formula of *p*\*, we show the following alternative:

- If *s_J_* < *s_A_* then 2*p*\* – 1 < 0 for all *m* ∈ (0,1) and *Div* is decreasing with respect of the maturation *m*.
- If *s_J_* > *s_A_* then there exists a threshold 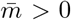 defined by 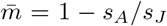 such that *Div* is maximal at 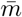 and if 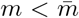, 2*p*\* – 1 > 0 and then *Div* is increasing with respect to *m* and for 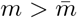, 2*p*\* – 1 < 0 and *Div* is decreasing with respect to *m*.

## Appendices

## A Properties of the demographic model in a single habitat

In this section, we describe the mathematical properties of the simple demographic model with two classes, the juveniles *J* and the adults *A*, in a single habitat. The dynamics of the population size **N**_*t*_ = (*J_t_*, *A_t_*) is given by

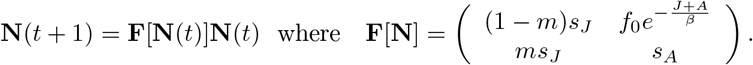

### Stationary state N^*^

The stationary states of (1) satisfy the following equation:

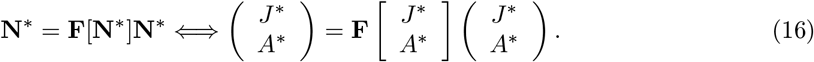

If we denote *N*^*^ = *J*^*^ + *A*^*^, then the system (16) is equivalent to:

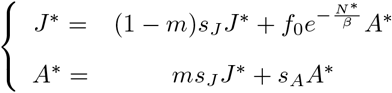

We deduce the expression *A*^*^ with respect to *J*^*^ and *N*^*^

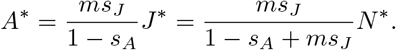

As a result we obtain:

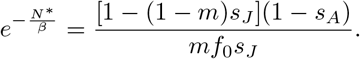

From hypothesis (*H*_3_), we know that *mf*_0_*s_j_*/[1 – (1 – *m*)*s_j_*](1 – *s_A_*) > 1 and

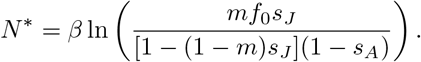

Finally we obtain the following expressions:

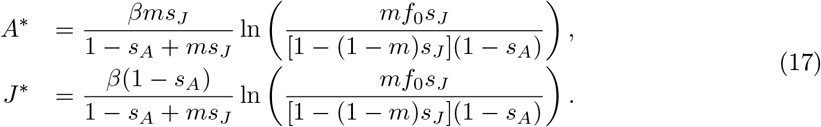

### Stability of N^*^ and Hopf bifurcation

#### Proposition 2.

*Let* **N**^*^ *be the stationary state defined by* (17). *Then* **N**^*^ *is stable if and only if f*_0_ < *f_c_ where f_c_ is defined by*:

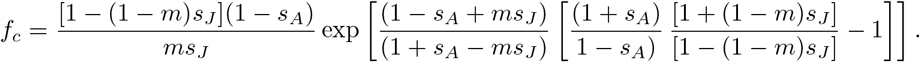

*Proof of Proposition 2*. We consider the stationary state **N**\* defined by (17). In order to determine the stability of this stationary state, we calculate the jacobian matrix **B** of **F** [**N**]**N** at point **N**\*:

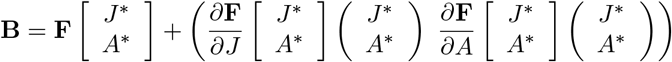

The partial derivatives of **F** are given by:

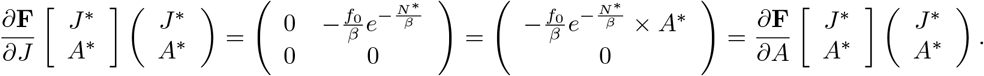

So we get:

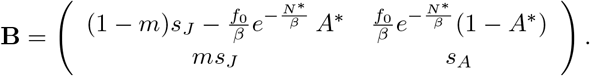

Local stability near the point of equilibrium **N**^*^ is given by the eigenvalues of the matrix **B**. Let *λ*_1_ and *λ*_2_ be the eigenvalues of the matrix **B**, we have det(**B**) = *λ*_1_λ_2_ and tr(**B**) = *λ*_1_ + *λ*_2_. More precisely, using the expressions of *N*^*^, we get:

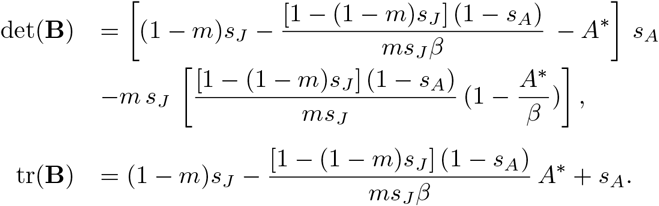

This state is stable if and only if |*λ*| < 1 for any eigenvalue *λ* of **B**.

- *λ* < 1 ⇔ λ – 1 < 0 and *λ* – 1 is an eigenvalue of (**B** – **I**). The eigenvalues of (**B** – **I**) are negative if and only if tr(**B** —**I**) < 0 and det(**B** – **I**) > 0. It is equivalent to

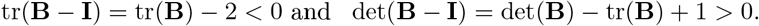
- *λ* > –1 ⇔ λ + 1 > 0 and *λ* + 1 is an eigenvalue of (**B** + **I**). The eigenvalues of (**B** + **I**) are positive if and only if tr(**B** + **I**) > 0 and det(**B** – **I**) > 0 and so:

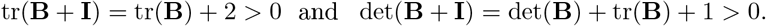

Hence the condition |λ| < 1 is equivalent to the following system of inequalities:

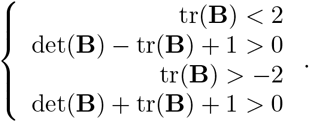

In the coordinate system (tr, det), our inequalities define the domain described in the figure 8. Let us look at the inequalities at the edge of this area.

- First we study the behaviour on (*A, B*) (see figure 8).

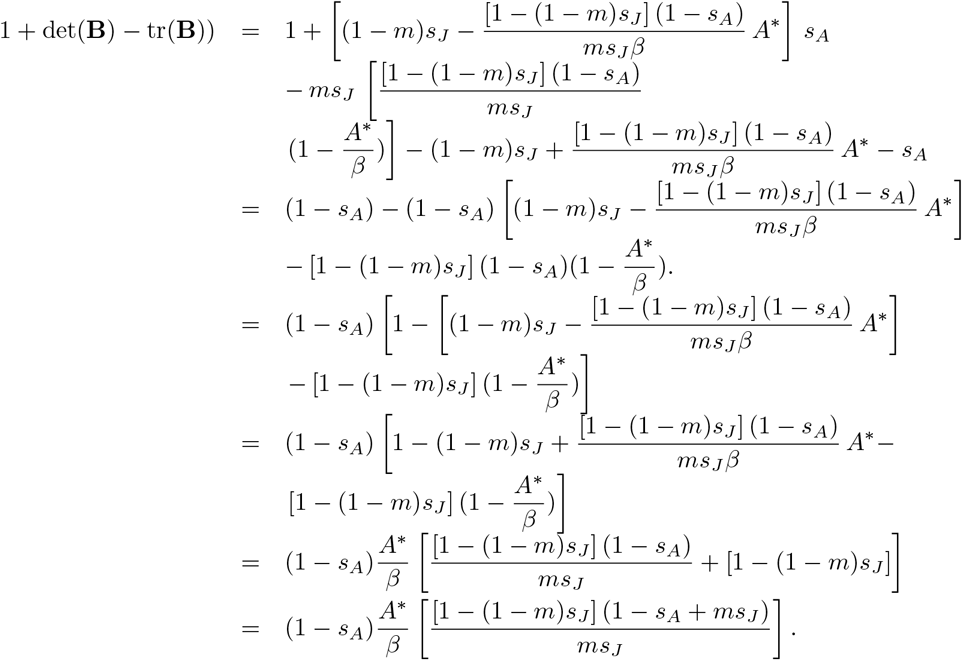 So, we always have 1 + det(**B**) – tr(**B**)) > 0 and thus, there is no bifurcation on this edge (*A, B*) of the stable area.
- Then we look at the condition tr(**B**) ≥ –2.

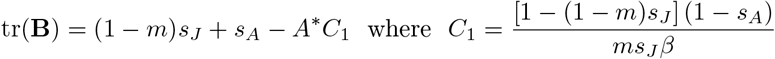 Thus, tr((**B**) > –2 if and only if *A*\**C*_1_ < 2 + (1 – *m*)*s_J_* + *s_A_*.
- On the other hand, det(**B**) = *A*\**C*_1_(1 – *s_A_* + *ms_J_*), from which we deduce:

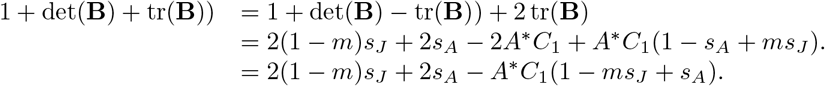 If 1 + det(**B**) + tr(**B**)) ≥ 0 then *A*\**C*_1_ ≥ 2. Thus we obtain *A*\**C*_1_ < 2 + (1 – *m*)*s_J_* + *s_A_* and we deduce tr(**B**) > –2.
- As a consequence, bifurcation may only occur only on (*C, A*) where 1 + det(**B**) + tr(**B**) = 0. From the previous computations, this equation is equivalent to *A*\**C*_1_(1 – *ms_J_* + *s_A_*) = 2[(1 – *m*)*s_J_* + *s_A_*]. And therefore we get:

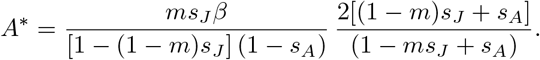 Now 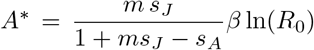 by definition, thus we obtain a condition *f*_0_ so that the dynamics of the habitat have bifurcations. In fact, we get:

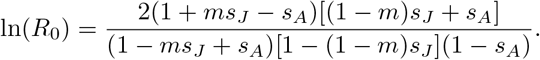 By combining this expression with the definition of R_0_,

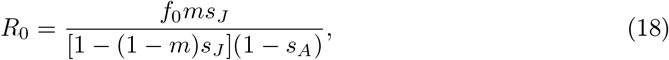

we get that:

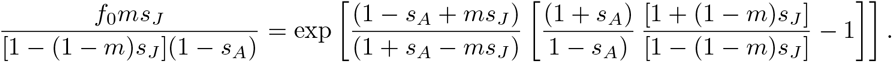 We deduce the value of intrinsic fertility *f_c_* such that the first bifurcation appears:

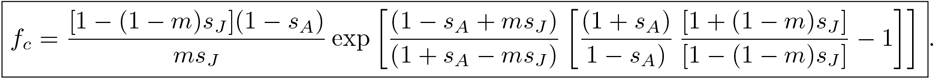

## B Mathematical properties of the asymptotic proportion *p*\*

### B.1 Extinction of a neutral fraction

From the Theorem 1, we know that the asymptotic proportion *p*\* is positive if the matrix **DF** [**N**\*] is primitive. However, if this property does not hold true, some alleles may eventually go extinct. For instance, in a metapopulation composed of 2 habitats in which migration only occurs from habitat 2 to habitat 1, that is **D**_42_=**0**, the projection matrix at equilibrium satisfies

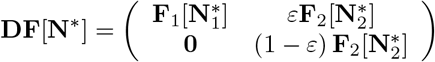

where **F**_*k*_ is defined by (7). It is obvious that **DF** [**N**\*] is not irreducible. Now assume that the allele **n** is initially only in habitat 1, that is **n**(0) = (**n**_1_(0), **0**). Then we get that

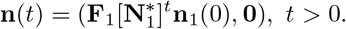

Now, let us show that the spectral radius of 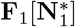 is less than 1. This is equivalent to prove that the net reproductive rate 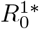 is less than 1 (Neubert and Caswell, 2000), where this quantity is defined by:

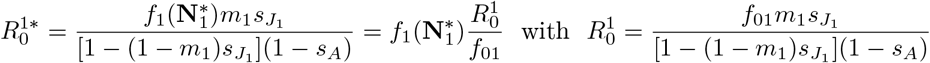

From the equation on **N**^*^ we can deduce the quantity 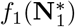. First we have

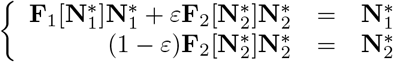

which provides the following relationship:

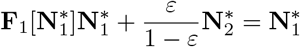

Developing this system we obtain:

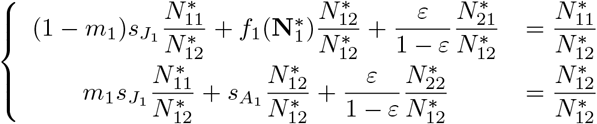

So, from the second equation of this system, we have:

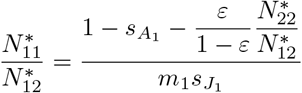

Therefore, we obtain:

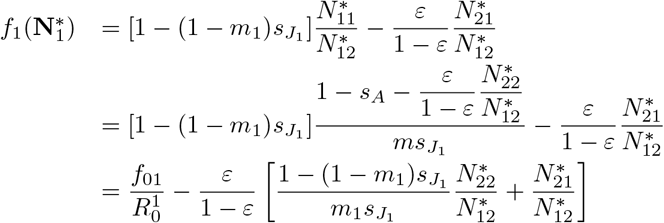

As consequences, we get

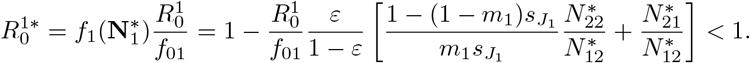

Finally, we conclude that **n**(*t*) converges toward **0** as t tends to ∞ which implies that *p*^*^ = 0 and the allele **n** goes extinct in the metapopulation.

### B.2 Asymptotic proportion *p*^*^ in time periodic steady state N^*^(*t*)

We investigate the case where the equilibrium **N**^*^(*t*) is a *T*–periodic steady state. We look at a neutral allele **n** which satisfies initially the following property:

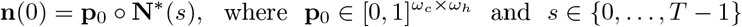

and ◦ is the Hadamard product. The vector **p**_0_ corresponds to the initial proportion of the neutral allele in the metapopulation. The initial configuration of the metapopulation is **N**\* (s) among the *T* possible configuration describes by the periodic steady state **N**\*. In this situation we have the following result

#### Proposition 3.

*Let* **N**\*(*t*) *be a T*–*periodic steady state of* (1) *and* **n** *be the solution of the following problem*

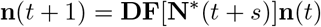

*starting from* **n**(0) = **p**_0_ ◦ **N**\*(*s*) *with* **p**_0_ ∈ [0,1]^*ω_c_* × *ω_h_*^ *and s* ∈ {0,…, *T* – 1}. *Then we have*

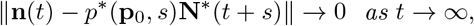

*where the asymptotic proportion p*^*^(**p**_0_,*s*) *is defined by*

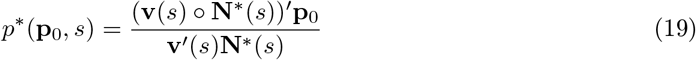

*where* **v**(*s*) *is the eigenvector associated to eigenvalue* 1 *of the transpose of the matrix* **C**(*s*) *defined by*

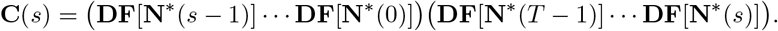

*Moreover, for any* (*t, s*) ∈ {0,…, *T* – 1} *with t* ≠ *s, the asymptotic proportions p*^*^(**p**_0_, *s*) *and p*^*^(**p**_0_,*t*) *are equal*, *p*^*^(**p**_0_,*s*) = *p*^*^(**p**_0_,*t*), *if and only if* 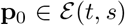 *where the set* 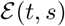 *is defined by*

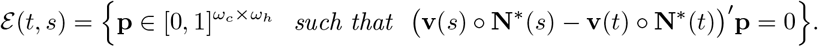

*The set* 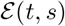 *is an hyperplan of codimension* 1 *which contains the vector* (1,…, 1) *or the whole space*.

The first part of this proposition is just a reformulation of Theorem 1. The second part shows that the asymptotic proportion *p*\* does depend on the initial configuration of the metapopulation **N**^*^. In particular, the asymptotic proportions *p*^*^(**p**_0_,*t*) and *p*^*^(**p**_0_,*s*) are not always equal if the dynamics of the metapopulation is such that **v**(*s*) ◦ **N**^*^(*s*) ≠ **v**(*t*) ◦ **N**\*(*t*). This difference explains why, in the time varying scenario, the diversity has multiple values. Still, we recover that if the initial proportion **p**_0_ has identical components then the asymptotic proportion are equal for any *s, t*.

*Proof of Proposition 3*. The first part of the Proposition is a reformulation of Theorem1 combined with the following equality

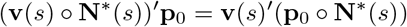

which is a consequence of the Hadamard product properties.

Now let us look at the second part of the proposition. From formula (19), it is straightforward to prove that *p*\*(**p**_0_,*s*) = *p*\*(**p**_0_,*l*) if and only if 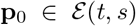. We just need to remark that **v**′(*s*)**N**\*(*s*) = **v**′(*l*) **N**\*(*l*) for all *s, l* because **v**(*s*) and **v**(*l*) are eigenvectors so we can choose them such that **v**′(*s*)**N**\*(*s*) = 1 = **v**′(*l*)**N**\*(*l*). From the definition of 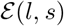 we know that it is an hyperplan of codimension at most 1. Moreover, it is of codimension 1 if and only if (**v**(*s*) ◦ **N**\*(*s*)) ≠(_v_(*t*)◦ **N**\*(*t*)).

Unfortunately, we are unable to characterize properly the equality **v**(*s*) ◦ **N**\*(*s*) = **v**(*t*) ◦ **N**\*(*t*). Numerically, we show that this equality does not occur for a large range of parameters of our model (see Fig. 9).

However, for the 2-periodic stationary state of the model with a single habitat we can prove that the set 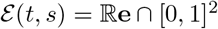 where **e** = (1,1).

*Proof of Proposition 3 in a single habitat*. In this proof, we consider the 2–periodic stationary state **N**\*(*t*) = (*J*(*t*), *A*(*t*)) of the following model

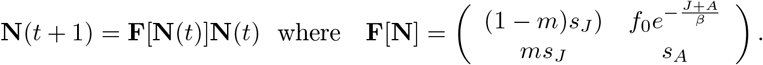

Our aim is to prove that **v**(0)◦**N**\*(0) ≠ **v**(1)◦**N**\*(1) where **v**(1) and **v**(0) are defined in Proposition3. Since 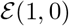 is an hyperplan orthogonal to the vector (**v**(0) ◦ **N**\*(0) – **v**(1) ◦ **N**\*(1)) and it contains the vector (1,1), this implies that the set 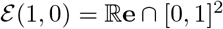.

To do so, let us first define **A**(*s*) as follows

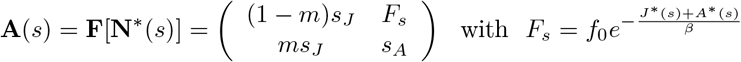

From the definition of **N**\*(*t*), we know that 1 is an eigenvalue of the matrix **C**(0) =**A**(1)**A**(0) because **N**\*(0) =**N**\*(2) = **A**(1)**A**(0)**N**\*(0). This implies that det(*I* –**C**(0)) = 0 where *I* is the identity matrix. This equality implies that *F*_1_ and *F*_0_ should satisfy the following equation

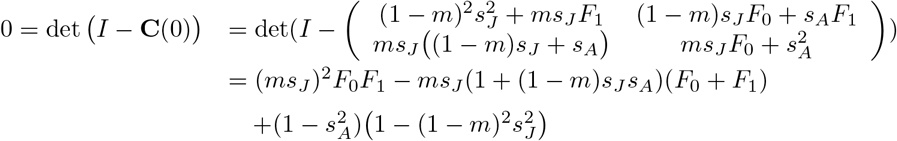

In particular, if we make the change of variable *Y*_0_ = *F*_0_ + *F*_1_ and *Y*_1_ = *F*_1_–*F*_0_, we get that *Y*_0_ and *Y*_1_ are on the hyperbole of the form

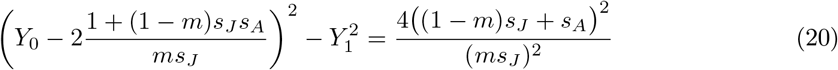

We can deduce that *F*_0_ and *F*_1_ are linked through this equation. Unfortunately, we are not able to compute at this stage *Y*_0_ or *Y*_1_.

Now let us look at the eigenvector **v**(0) of (**C**(0))′ associated to eigenvalue 1 and such that **v**(0)′**N**\*(0) = 1. Thus, we have

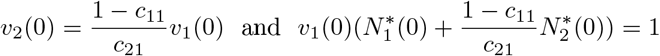

Since **N**\* (0) is an eigenvector of **C**(0) associated to the eigenvalue 1, we get that 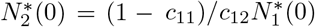. Thus we get that

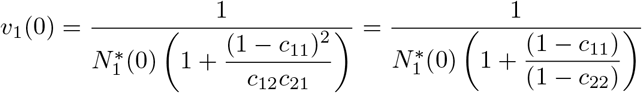

because det(*I* – **C**(0)) = (1 – *c*_11_)(1 – *c*_22_) – *c*_12_*c*_21_ = 0. Let us now compute **v**(0) ◡ **N**\*(0).

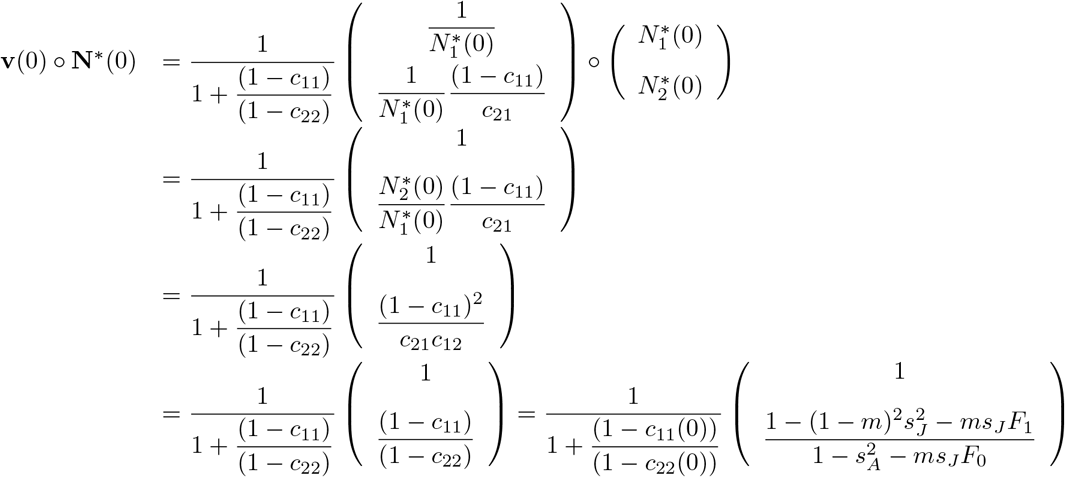

Similarly, we can compute **v**(1) ◦ **N**^*^(1) as follows

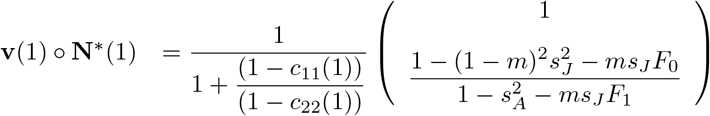

Thus we can see that **v**(0) ◦ **N**\*(0) = **v**(1) ◦ **N**\*(1) if and only if

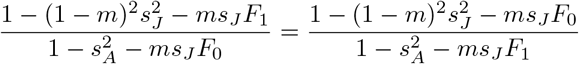

Thus, we have either *F*_1_ = *F*_0_ or 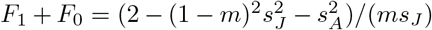. From the equation (20), we deduce that 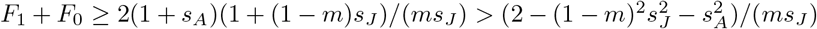. Thus, we get *F*_0_ = *F*_1_. However, *F*_0_ = *F*_1_ because **N**^*^(*t*) is a 2-periodic stationary state (**N**\*(0) ≠ **N**\*(1)) and **A**(0) ≠ **A**(1), otherwise **N**\*(1) = **N**\*(0).

In conclusion, we get that **v**(0) ◦ **N**^*^(0) ≠ **v**(1) ◦ **N**^*^(1) and thus the set 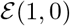 is of codimension 1 and since **e** = (1,1) is in 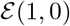, we have 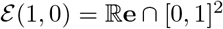.

We run some simulations to show that the asymptotic proportion *p*\* can take different value when the steady state becomes periodic (see Fig. 9).

### B.3 Speed of convergence in a single habitat

We consider the stationary state **N**^*^ of the model in a single habitat with two stages which is defined by (17). Then, any allele **n** starting from **n**(0) satisfies the following linear equation

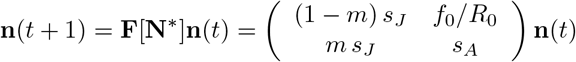

with *R*_0_ defined by (18). Then the reproduction matrix **F**[**N**^*^] is defined by:

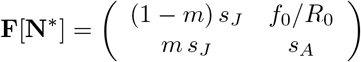

The matrix **F**[**N**\*] is diagonalisable with eigenvalues 1 and |λ_2_| < 1 defined by

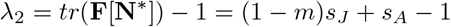

Thus, the allele **n**(*t*) can be decomposed as follows

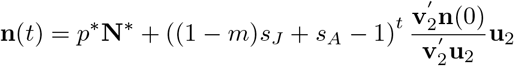

where **N**\* is the stationary state and *p*\* is defined by the Theorem 1, and **u**_2_ and **v**_2_ are respectively the right and the left eigenvector of the matrix **F**[**N**\*] associated to the eigenvalue λ_2_. A direct computation shows that

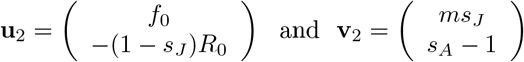

Moreover, we know that |λ_2_| < 1, however, λ_2_ might be negative. In particular, if 0 < (1 – *m*)*s_J_* + *s_A_* – 1 < 1, the allele **n**(*t*) converge monotonically towards *p*\***N**\* (see Fig. 10(a)). While if –1 < (1 – *m*)*s_J_* + *s_A_* – 1 < 0, the allele **n**(*t*) will converge with damped oscillation around the limit *p*^*^**N**^*^ (see Fig. 10(b)).

## C The individual–based model of neutral genetic fractions

We consider a model of a metapopulation of size *N_s_* and composed of 2 habitats and 2 stages in each habitat. Let **X**_*t*_ = (*J*_*t*, 1_, *A*_*t*, 1_, *J*_*t*, 2_, *A*_*t*, 2_) where *J*_t_, *i* is the number of juveniles in each habitat *i* and *A*_*t, i*_ is the number of adults in each habitat *i*. For each habitat *i*, adults give birth according to a Poisson law of parameter *f_i_*(*J_t, i_*/*N_s_*, *A_t, i_*/*N_s_*) and they die at a rate *s_Ai_* according to a Bernoulli law. Similarly, for each habitat *i*, juveniles survive at rate (1 – *m_i_*)*s_Ji_* according to a Bernoulli law. And each individual migrates from habitat *i* to habitat *j* with a rate 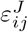 or 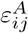 depending on whether he is juvenile or adult.

We suppose that the evolution of this metapopulation is iterative in discrete time and we obtain the stochastic model as follows:

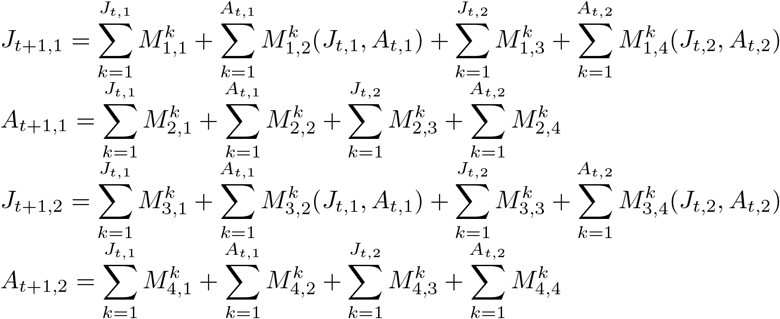

- where 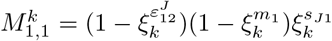 and the *ξ_k_* are Bernoulli i.i.d variables with parameter respectively 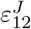, *m*_1_ and *s*_*J*1_.
- where 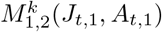 is a sum of *F*^1^(*J*_*t*,1_, *A*_*t*,1_) Bernoulli variables of parameter 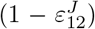 where *F*^1^(*J*_*t*, 1_, *A*_*t*, 1_) is a Poisson variable with parameter *f*_1_(*J*_>*t*, 1_/*N*_*s*_, *A*_*t*, 1_/*N_s_*).
- where 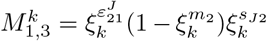 and the *ξ_k_* are Bernoulli i.i.d variable with parameter respectively 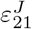, *m*_2_ and *s*_*J*2_.
- where 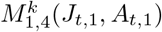 is a sum of *F*^2^(*J*_*t*, 2_, *A*_*t*, 2_) Bernoulli variables of parameter 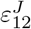 where *F*^2^(*J*_*t*, 2_, *A*_*t*, 2_) is a Poisson variable with parameter *f*_2_(*J*_*t*, 2_/*N_s_*, *A*_*t*, 21_/*N_s_*). In fact, 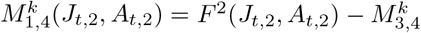.
- where 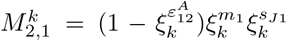 and the *ξ_k_* are Bernoulli i.i.d variables with parameters respectively 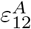, *m*_1_ and *s*_*J*1_.
- where 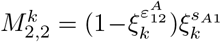 and the *ξ_k_* are Bernoulli i.i.d variables with parameter respectively 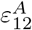 and *s*_*J*1_.
- where 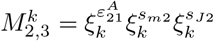 and the *ξ_k_* are Bernoulli i.i.d variables with parameter respectively 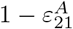, *m*_2_ and *s*_*j*2_.
- where 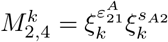 and the *ξ_k_* are Bernoulli i.i.d variables with parameter respectively 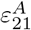 and *s*_*A*2_.
- where 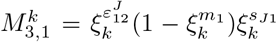 and the *ξ_k_* are Bernoulli i.i.d variables with parameter respectively 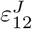, *m*_1_ and *s*_*J* 1_
- where 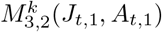 is a sum of *F*^1^(*J*_*t*,1_, *A*_*t*, 1_) Bernoulli variables of parameter 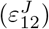 where *F*^1^(*J*_*t*, 1_, *A*_*t*, 1_) is a Poisson variable with parameter *f*_1_(*J*_*t*, 1_/*N_s_*, *A*_*t*, 1_/*N_s_*). In fact, 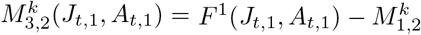.
- where 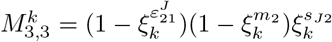 and the *ξ_k_* are Bernoulli i.i.d variables with parameter respectively 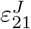, *m*_2_ and *s*_*J*2_.
- where 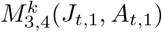 is a sum of *F*^2^(*J*_*t*, 2_, *A*_*t*, 2_) Bernoulli variables of parameter 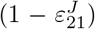 where *F*^2^(*J*_*t*, 2_, *A*_*t*, 2_) is a Poisson variable with parameter *f*_2_(*J*_*t*, 2_/*N_s_*, *A*_*t*, 21_/*N_s_*).
- where 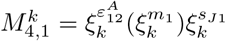 and the *ξ_k_* are Bernoulli i.i.d variables with parameter respectively 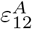, *m*_1_ and *s*_*J*1_.
- where 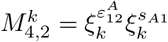 and the *ξ_k_* are Bernoulli i.i.d variables with parameter respectively 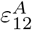 and *s*_*A*1_.
- where 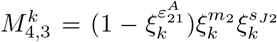 and the *ξ_k_* are Bernoulli i.i.d variables with parameter respectively 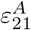, *m*_2_ and *s*_*J*2_.
- where 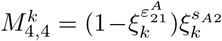 and the *ξ_k_* are Bernoulli i.i.d variables with parameter respectively 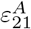 and *s*_*A*2_.

So, we have:

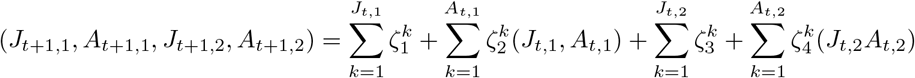

with 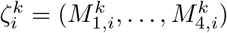. We have:

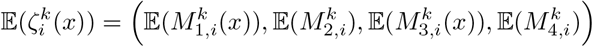

So, we obtain:

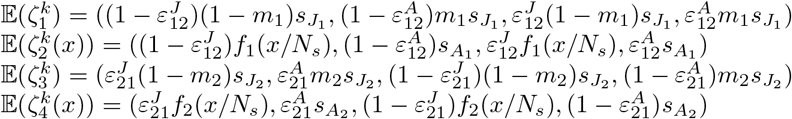

So we can see that 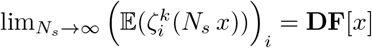

Likewise, if we consider the covariance matrix 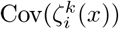 for *i* ∈ {1,…, 4}, according to the independence of all the variables we obtain:

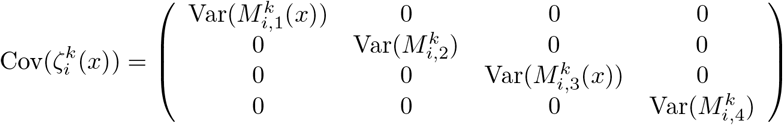

where the only dependence in *x* is for Var(*M*_1,2_(*x*)) and Var(*M*_3,4_(*x*)) which are equals respectively to

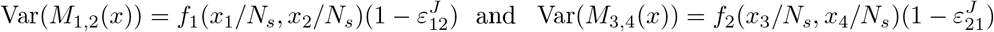

We obtain that 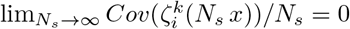 for any *i*. Now we can apply the result of (Adam,2016) and thus the stochastic model converges in probability when the size *N_s_* tends to + ∞ to our deterministic model that we have considered above, more precisely we have:

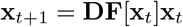

where

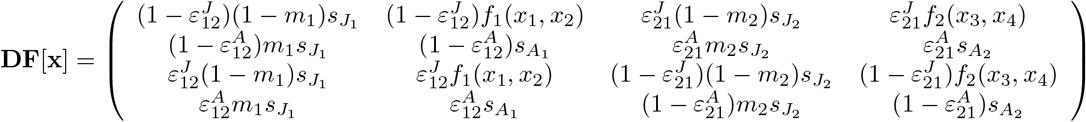

and x_*t*_ = (*J_1,t_*, *A_1,t_*, *J_2,t_*, *A_2,t_*).

### C.1 Numerical simulations

Our stochastic model converges when the size of the population becomes infinite towards our deterministic model (1). We illustrate this convergence and the different behaviour of the stochastic model in the following numerical simulations.

First, we compare the outcome of the stochastic model described in Fig.2 with different population size. In Fig.11, we see that the stochastic model can experience genetic drift when the population size is low *N_e_* ≈5.10^2^(*N_s_* = 5), while genetic drift is lees likely to occur when the population size increases (see *N_e_* ≈ 5.10^5^(*N_s_* = 5.10^3^).

Similarly, we see from Fig.12 that on average the behaviour of the stochastic model converges to the behaviour of the deterministic model even if genetic drift may occur. In particular, we see that the variance among trajectories vanishes when the population becomes large enough, which shows that genetic drift is unlikely to occur in this situation.

